# MBNL1 and RBFOX1 co-regulate alternative splicing events transcriptome-wide through a conserved buffering mechanism

**DOI:** 10.1101/2021.04.20.440659

**Authors:** Melissa Hale, Joseph Ellis, Ryan Meng, Sunny McDaniel, Amy Mahady, Stacey Wagner, Jared Richardson, Ona McConnell, Eric Wang, J. Andrew Berglund

## Abstract

Alternative splicing (AS) is controlled by *cis*-regulatory elements recognized by networks of *trans*-acting factors. Here we investigate modes and mechanisms of AS co-regulation by MBNL1 and RBFOX1, two RNA binding proteins (RBPs) critical for developmental AS transitions. We generated two cell models that express each RBP under separate inducible promoters. Transcriptome-wide categorization of the impacts of RBFOX1 expression on MBNL1 splicing revealed a common co-regulatory mode through which RBFOX1 *buffers* MBNL1 dose-dependent splicing regulation by reducing the total range of exon inclusion or exclusion. Minigene mutational analysis and *in vitro* binding experiments suggest that this *buffering* mechanism occurs through a shared *cis*-regulatory element previously unidentified as critical for MBNL1-dependent activity. Overall, our studies define a conserved co-regulatory mechanism through which RBFOX1 and MBNL1 can fine-tune and provide redundancy for AS outcomes. These studies indicate overlapping use of RNA motifs with potential implications for when activity of RBPs is disrupted.

## Introduction

Alternative splicing (AS) is an intricate and dynamic step of RNA processing whereby a single gene can produce multiple RNA isoforms that dramatically impact the complexity of the cellular transcriptome and proteome. Transcriptomic studies have shown that 90-95% of multi-exon genes produce alternatively spliced transcripts^1–3^. AS is regulated through an interaction network of *cis-*regulatory sequence elements within pre-mRNA transcripts and their recognition by *trans-*acting factors, most-notably RNA binding proteins (RBPs). General trends in splicing regulation have been established in the context of specific *cis*-regulatory motifs utilized by individual RBPs. However, the deeper fundamental rules that govern accurate and coordinated use of the “splicing code” by groups of RBPs to facilitate proper cellular differentiation and tissue development remains poorly understood^4–7^. This lack of understanding is further exaggerated when examining how overall changes in cellular RBP concentrations and aberrant AS splicing outcomes can have profound impacts in the context of disease. Alterations in functional RBP levels have been linked to multiple classes of diseases including cancer, heart disease, and neurodegenerative disorders^8–11^. For the latter, one of the best examples is myotonic dystrophy (DM) which results from expression of an expanded CUG (DM type 1) or CCUG (DM type 2) RNA. These repeat RNAs serve as a sink for muscleblind-like (MBNL) proteins; a family of RBPs that regulate fetal to adult AS transitions critical for proper muscle, heart, and central nervous system (CNS) development^12,13^. Reduction of functional MBNL protein levels via this toxic RNA gain-of-function mechanism leads to global perturbations of AS established as causative of a number of DM disease symptoms^14–16^.

MBNL proteins have been shown to bind to YGCY *cis-*regulatory motifs to regulate AS outcomes of target pre-mRNAs in a positional-dependent manner whereby cassette exon inclusion is generally promoted through binding to downstream RNA motifs and repressed through use of upstream intronic elements^17^. Previous studies have indicated that MBNL-dependent AS responds to changes in MBNL concentrations across a broad, dynamic range of expression^18,19^. Using a titratable MBNL1 expression system to characterize the behavior of individual MBNL1-regulated AS events, it was previously reported that different amounts of MBNL are required to achieve half-maximal regulation with variable steepness of the dose-response curves^18^. While several studies have focused on the characterization of MBNL-dependent splicing regulation^13^, less work has been done to identify and characterize the role of other splicing factors that may coordinate with or antagonize MBNL splicing activity with the exception of CELF1^20–22^. Recent reports have indicated that MBNL and another family of RBPs, RBFOX, co-regulate a subset of AS events, including some events that are mis-regulated in DM^23,24^. RBFOX proteins regulate AS in the same positional-dependent manner as MBNL proteins via binding to (U)GCAUG motifs within target RNAs with high affinity and specificity^25–27^. Like the family of MBNL proteins, RBFOX protein expression is critical for proper skeletal muscle, heart, and CNS development^28–30^. Additionally, it was previously demonstrated that MBNL and RBFOX proteins bind to the expanded CCUG RNA repeats found in DM2, suggesting that these two splicing factors have the capacity to recognize similar RNA motifs^31^. Collectively, the similarities in mechanisms of AS regulation, tissue-specific expression, overlap of regulated events, and the potential recognition of the same or similar RNA regulatory sequences suggests that unidentified and under-appreciated mechanisms of AS co-regulation shared by these two RBPs may exist.

To evaluate possible mechanisms of AS co-regulation we utilized two cell lines in which the concentrations of RBFOX1 and MBNL1 could be systematically and independently expressed in a dose-dependent manner. Human embryonic kidney (HEK-293) and mouse embryonic fibroblast (MEF) cell lines were engineered with titratable expression of RBFOX1 and MBNL1 to identify large populations of AS events regulated by one or both factors. Categorization of splicing events identified via RNAseq into predicted co-regulatory groups revealed that RBFOX1 affects MBNL1 dose-dependent splicing transcriptome wide. Specifically, we found a high number of events in which RBFOX1 expression reduces the overall response of target AS events to changes in MBNL1 cellular concentrations without impacting maximal exon inclusion levels. We propose that this previously unidentified *buffering* co-regulatory mechanism is facilitated through shared use of a UGCAUG *cis*-regulatory element that contains a suboptimal MBNL1 motif (UGCA). Overall, the experimental observations using these dosing cell lines define a complex, conserved, and interactive network of AS regulation shared by MBNL1 and RBFOX1 that provides precise control to modulate AS outcomes.

## Results

### RBFOX1 expression buffers MBNL1 dose-dependent splicing regulation of INSR minigene splicing reporter

To evaluate the ability of MBNL1 and RBFOX1 to co-regulate alternative splicing outcomes we assayed if co-expression of both RBPs impacted regulation of the *INSR* exon 11 AS event via a minigene reporter system^32^. The *INSR* minigene contains a single, putative UGCAUG RBFOX binding site embedded within several YGCY RNA elements in intron 11, many previously identified as critical for MBNL-dependent exon 11 inclusion^32–35^ (Fig 1a). RBFOX1 has previously also been shown to independently promote *INSR* exon 11 inclusion^31^. We assayed exon 11 inclusion using a previously generated tetracycline-inducible HA-MBNL1 HEK-293 cell line transiently transfected with a HA-RBFOX1 expression vector. Expression of HA-MBNL1 alone via doxycycline treatment following transfection of the minigene reporter led to significantly increased *INSR* exon 11 inclusion as quantified by *percent spliced in* (Ψ) (i.e. percent exon inclusion) (Fig 1b). A similar, albeit reduced level, of inclusion was observed following expression of HA-RBFOX1 in the absence of MBNL1 induction (Fig 1b). While co-expression of both splicing factors further enhanced exon inclusion above that of each RBP individually, the overall ΔΨ was not additive as maximal exon inclusion (i.e. Ψ = 1.0) was not achieved (Fig 1b). We observed a similar pattern of splicing activity for the MBNL1-dependent *Nfix* exon 8 minigene reporter^36^ where expression of either RBP alone induced exon exclusion while co-expression led to a small, but not additive combinatorial effect (Fig S1). The lack of an additive impact on exon inclusion under co-expression of RBFOX1 and MBNL1 suggests that these two RBPs likely do not function independently but rather in a cooperative manner to modulate exon inclusion.

**Figure 1:**
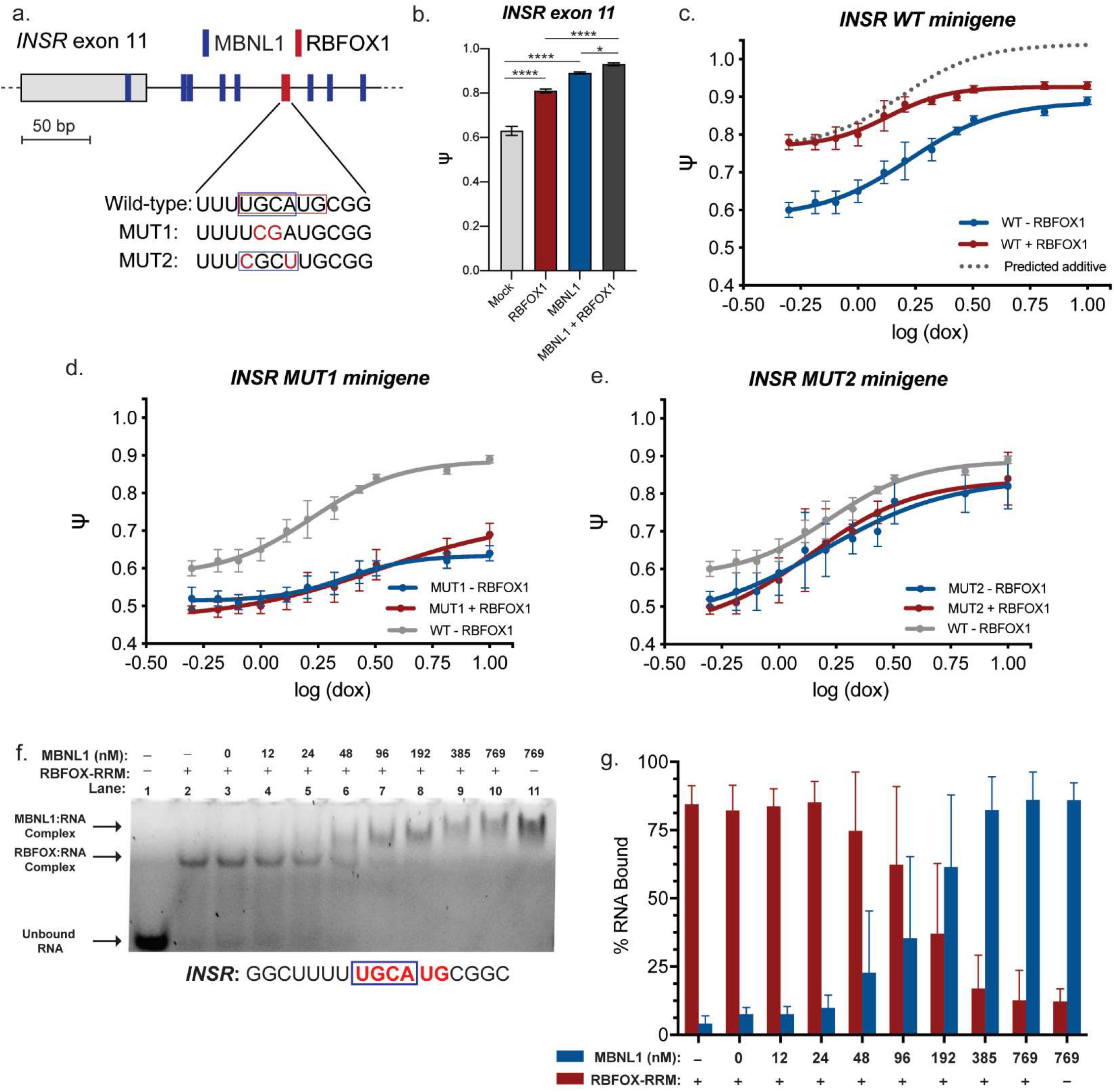
RBFOX1 buffers MBNL1 dose-dependent splicing regulation of INSR exon 11 minigene through binding to a shared cis-regulatory element. (a) Distribution of MBNL1 (YGCY) and RBFOX1 (UGCAUG) binding motifs in wild-type, MUT1, and MUT2 INSR exon 11 minigenes. (b) Bar-plot representation of cell-based splicing assay with wild-type INSR exon 11 minigene in HA-MBNL1 tetracycline-inducible HEK-293 cells18 transfected with either empty vector or HA-RBFOX1 individually or in combination. HA-MBNL1 expression was induced via doxycycline (10 ng/mL). Data represented as mean percent exon inclusion (Ψ) ± SEM, n=3. (* p < 0.05, **** p < 0.0001, one-way ANOVA). (c-e) MBNL1 dose-response curves for wild-type (WT), MUT1, and MUT2 INSR minigenes in presence or absence of HA-RBFOX1 expression. Ψ was measured at each concentration and plotted as a function of log (dox) and fit to a four-parameter dose-curve. Data represented as mean ± SEM, n = 3. Representative native gels and quantitative parameters derived from dose-response curves are supplemented (Fig S3, Table S1). (f) Representative competitive EMSA showing binding of both purified MBNL1 and RBFOX-RRM to a model RNA containing the INSR intron 11 putative UGCAUG RNA motif and surrounding sequence. (g) Quantification of percent RNA bound by either RBFOX1 or MBNL1 upon challenge with increasing concentrations of MBNL1 (n = 3).

To further discern the effects of RBFOX1 and MBNL1 co-expression on AS of the *INSR* minigene, we measured changes of exon 11 inclusion across a range of MBNL1 protein expression in the presence or absence of HA-RBFOX1. Following transfection of a HA-RBFOX1 expression vector, MBNL1 expression was induced by titration of doxycycline to produce a range of MBNL1 concentration and splicing activity (Ψ) plotted as a function of log (doxycycline) (i.e. log(dox)), a proxy for HA-MBNL1 protein concentration)^18^. These data were fit to a four-parameter dose curve to derive quantitative parameters that describe the observed patterns of splicing regulation in either the presence or absence of RBFOX1 expression (Fig 1c), including EC_50_, slope, and span (ΔΨ)^18^. The slope of the response curve provides a relative measure of cooperativity while the EC_50_ value indicates how much protein is required to obtain splicing regulation at 50% of maximum Ψ (Fig S2).

Co-expression of HA-RBFOX1 led to an overall reduction in ΔΨ covered over the MBNL1 concentration gradient (ΔΨ_*-RBFOX1*_ = 0.26 vs. ΔΨ_*+RBFOX1*_ of 0.12, Fig 1c). Interestingly, the observed change in Ψ occurred in a non-uniform manner at each MBNL1 dose. While at lowest levels of HA-MBNL co-expression of HA-RBFOX1 significantly increased *INSR* exon 11 inclusion (ΔΨ_*LOW*_ = 0.18), induction of higher cellular concentrations of MBNL1 resulted in only a minimal increase in Ψ (ΔΨ_*HIGH*_ = 0.04). Based on the inclusion level at the lowest MBNL1 expression level, a truly additive change (Fig 1c, grey curve) would be expected to exceed the maximum level of exon inclusion. However, the observed pattern of AS co-regulation does not appear to be due to an independent, combinatorial effect of both RBPs. Instead, MBNL1 and RBFOX1 appear to cooperatively interact to facilitate a co-regulatory model that we have termed *buffering*. More specifically, RBFOX1 reduces the overall ΔΨ of the MBNL1 dose-response by significantly affecting Ψ at low MBNL1 levels with minimal impacts at high MBNL1 concentrations. This conclusion is supported by the fact that almost half the amount of MBNL1 is required to affect an equivalent amount of splicing regulation when HA-RBFOX1 is co-expressed (log (EC_50_) = 0.22 ± 0.03 (+ RBFOX1) vs. 0.13 ± 0.03 (- RBFOX1)). Overall, MBNL1 and RBFOX1 appear to operate through a shared mechanism of AS regulation within the context of the *INSR* exon 11 minigene splicing reporter.

### MBNL1 and RBFOX1 co-regulate INSR exon 11 inclusion though a shared RNA cis-regulatory element

To understand the mechanism through which MBNL1 and RBFOX1 regulate *INSR* exon 11 inclusion, we tested the function of the predicted UGCAUG RBFOX binding site within the downstream intron 11. This *cis*-regulatory motif was mutated to U<u>CG</u>AUG to generate the *INSR* MUT1 minigene (Fig 1a). Inversion of this GC dinucleotide has been previously shown to abrogate RBFOX1 binding and prevent AS regulation^37–39^. As predicted, co-expression of HA-RBFOX1 had no effect on splicing of the *INSR* MUT1 minigene (Fig 1d, red curve). Surprisingly, this mutation significantly blunted MBNL1 dose-dependent splicing regulation compared to WT *INSR* (Fig 1d, compare grey and blue curves). While this result is consistent with RBFOX1 regulating exon inclusion through this specific *cis-*regulatory element, the substantial impact on MBNL1 splicing regulation was unexpected given that (1) many MBNL1 YGCY binding motifs are present within intron 11 and have previously been shown facilitate MBNL1-dependent exon 11 inclusion^32–35^ and (2) the RBFOX UGCAUG binding motif does not contain what is considered a MBNL1 consensus sequence^17^, although <u>UGCA</u> has been identified as a low affinity MBNL1 binding motif^27^. These results suggested that both RBPs use this single site to regulate exon 11 inclusion despite it being a non-optimal MBNL1 binding motif.

To validate if MBNL1 dose-dependent splicing regulation of *INSR* exon 11 is contingent on use of the UGCAUG RNA motif we generated a minigene in which the sequence was mutated to <u>C</u>GC<u>U</u>UG (*INSR* MUT2 minigene) (Fig 1a). This mutation disrupts the RBFOX binding site as in *INSR* MUT1 but generates a new, YGCY MBNL binding site in the same position as in the WT reporter (CGCU). Alteration of this motif nearly fully restored MBNL1’s ability to regulate splicing and, as predicted, co-expression of HA-RBFOX1 had no impact on AS (Fig 1e). Characterization of the MBNL1 dose-response for all three *INSR* minigene variants in the presence or absence HA-RBFOX1 expression are consistent with a model in which MBNL1 and RBFOX1 bind an overlapping site to modulate exon 11 inclusion. Furthermore, these experiments support a *buffering* co-regulatory mechanism whereby at low concentrations of MBNL1, RBFOX1 binds to the target site to promote exon inclusion. As the cellular concentrations of MBNL1 protein increases, MBNL1 may outcompete RBFOX1 to bind the same, overlapping site such that the maximum Ψ is achieved.

To evaluate this proposed model, we conducted a competitive electrophoretic mobility shift assay (EMSA) using a section of *INSR* intron 11 containing the putative RBFOX UGCAUG *cis-*regulatory element; no YGCY motifs were included in this model RNA sequence (Fig 1f). This RNA probe was pre-equilibrated with a sustained concentration (192 nM) of purified RBFOX RNA recognition motif (RRM) and then challenged with increasing concentrations of purified, GST-tagged MBNL1 (Fig 1f). When MBNL1 and RBFOX-RRM were equilibrated alone nearly all RNA (~85%, Fig 1g) was bound by either RBP independently. Furthermore, MBNL1 and RBFOX-RRM alone produced a visually distinct shift compared to free RNA probe and each other (Fig 1f, lanes 1, 2, and 11). When both MBNL1 and RBFOX-RRM were present (Fig 1f, lanes 4-10), as the concentration of MBNL1 increased a shift away from the RBFOX-RRM complex toward the MBNL1-RNA complex was detected. At equivalent protein concentrations (Fig 1f, lane 8), the RBFOX-RNA complex was barely visible and a strong MBNL1-RNA complex was detected. Percentage of RNA bound in each lane containing either RBFOX-RRM or MBNL1 was quantified (Fig 1g). These combined observations along with the lack of any clear or distinct ternary complexes across the titration of MBNL1 protein suggests independent binding of each RBP to the RNA probe. Collectively, results from minigene reporter splicing assays and EMSAs indicate that RBFOX1 and MBNL1 co-regulate *INSR* exon 11 inclusion through the use of a shared *cis*-regulatory motif traditionally classified as a specific, RBFOX1 binding element. Mechanistic insights gained through these studies indicate that these two RBPs may have the potential to co-regulate other splicing events through shared RNA regulatory regions.

### Dual-inducible RBFOX1 & MBNL1 cellular expression models allow for independent and precise control of RBP concentration to evaluate AS co-regulation

Protein expression levels using transient transfection of a plasmid, such as expression of HA-RBFOX1 within our *INSR* minigene experiments (Fig 1), is generally less robust due to the effects of experimental timing and transfection efficiency. For this reason, to further explore MBNL1 and RBFOX1 AS co-regulation, we choose to generate two distinct inducible cellular expression systems to more precisely and independently control levels of both RBPs into both mouse and human cell lines. Both a doxycycline-inducible MBNL1 and ponasterone A (ponA) - inducible RBFOX1 expression construct were stably integrated into HEK-293 and *Mbnl1*^−/−^ / *Mbnl2*^−/−^ mouse embryonic fibroblasts (MEFs)^40^. While the same mOrange-tagged RBFOX1 expression construct was integrated into both cell lines, the HEK-293 cells contained a previously integrated HA-MBNL1 construct^18^ while a GFP-tagged MBNL1 expression construct was used for the MEFs (Fig 2a). Within the context of these modified cell lines, hereafter referred to as double dosing HEKs (ddHEKs) or double dosing MEFs (ddMEFs), the use of doxycycline and/or ponA can be used to independently and precisely control expression of MBNL1 and RBFOX1, respectively.

**Figure 2:**
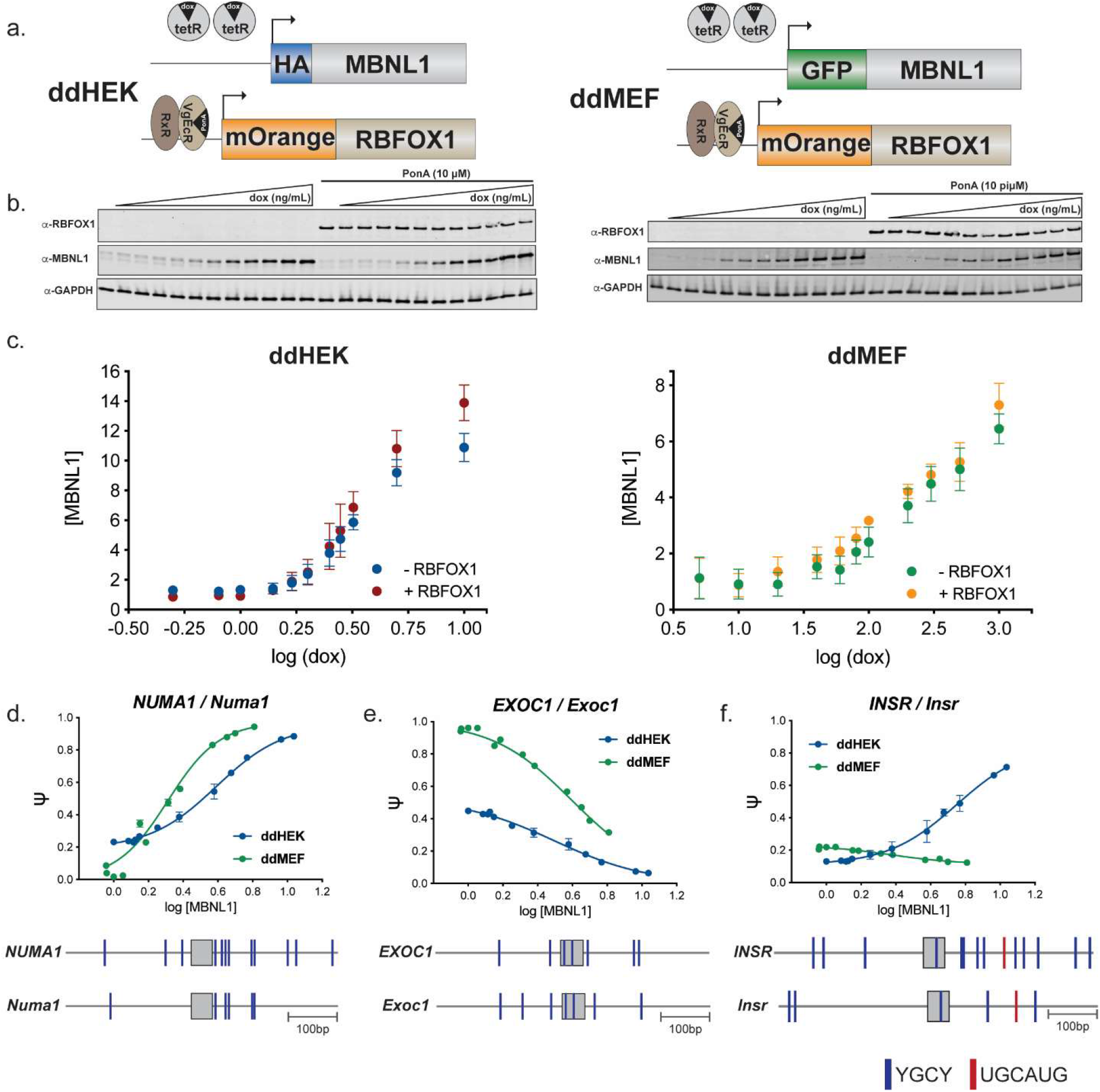
MBNL1 dose-dependent splicing is broadly conserved in novel ddHEK and ddMEF double dosing cell lines in which MBNL1 and RBFOX1 expression levels are independently and precisely controlled. (a) Model depicting inducible MBNL1 and RBFOX1 constructs in HEK-293 cells (ddHEKs) and Mbnl1^−/−^/Mbnl2^−/−^ MEFs (ddMEFs). HA-MBNL1 (ddHEK) or GFP-MBNL1 (ddMEF) expression is controlled via a tetracycline-inducible system. Doxycycline (dox) treatment induces expression of the full-length MBNL1 mRNA by de-repressing inhibition of transcription by binding to the tet repressor (tetR). Independently, expression of mOrange-tagged RBFOX1 is controlled via an ecdysone-inducible system. (b) Representative immunoblot showing gradient of MBNL1 expression produced via titration of doxycycline in the presence or absence of RBFOX1 expression induced via ponasterone A treatment. (c) Quantification of MBNL1 immunoblot normalized to GAPDH loading control in the presence and absence of RBFOX1 expression plotted as a function of log (dox). Data represented as mean ± SEM, n = 4. (d-f) MBNL1 dose-response curves for orthologous cassette exons NUMA1/Numa1, EXOC1/Exoc1, and INSR/Insr in ddHEK and ddMEF cell lines. Data represented as mean Ψ ± SEM (n = 3 - 5) plotted against log [MBNL1] levels as quantified by immunoblot (Fig 2c). Quantitative parameters derived from MBNL1 dose-response are available in Fig S4b, d. Distribution of YGCY motifs within 250-300bp upstream and downstream of each orthologous cassette exon is also displayed. In the case of INSR/Insr, the distribution of the RBFOX1 UGCAUG binding motif is also annotated.

For both cell lines, in the absence of dox or ponA, minimal levels of the integrated MBNL1 or RBFOX1 expression are detected by immunoblot (Fig 2b). Treatment with ponA induces high levels of RBFOX1 protein expression in both lines (Fig 2b). Similar to previous reports^18^, the titration of doxycycline in our cells leads to a broad range of a HA-MBNL1 or GFP-MBNL1 expression that can be precisely controlled (0-10 ng/mL for ddHEK, 0-1000 ng/mL for ddMEF) (Fig 2b). Quantification of MBNL1 protein levels via immunoblot revealed a 6 to 10 fold increase in maximal MBNL1 cellular levels in human and mouse cell lines, respectively (Fig 2c). MBNL1 expression levels were not significantly altered upon expression of RBFOX1 in either dosing model (Fig 2c). These two cell models provide the ability to alter the expression of either RBP alone or in combination in order to compare AS co-regulation from two distinct species.

### MBNL1 dose-dependent splicing regulation is broadly preserved in both human and mouse dosing cell models

Using our double dosing cell models we first chose to validated MBNL1 dose-dependent splicing independent of RBFOX1 co-expression. While previous work has identified and characterized MBNL1 dose-dependent splicing regulation^18,19^, little work has been performed to date to assess if concentration-dependent splicing outcomes are conserved across species. We selected several endogenously expressed cassette exons previously shown to be impacted by MBNL1 expression^41–43^, including *INSR*/*Insr* exon 11, and quantified exon inclusion levels across the MBNL1 concentration gradient in both systems. Ψ values were then plotted against log [MBNL1] for both MBNL1-regulated inclusion and exclusion events (Fig 2d-f and Fig S4). Consistent with previous studies, estimated EC_50_ values for cassette exon events assayed within the ddHEKs indicated that a range of MBNL1 concentrations are required to effectively regulate exon inclusion with variable magnitudes of cooperativity as measured by slope of the dose-response^18^ (Fig S4a-b). These same general trends were also observed within the ddMEFs (Fig S4c-d). In fact, several orthologous cassette exons expressed in both systems demonstrated similar regulation as quantified by EC_50_ and slope upon MBNL1 titration, possibly as a consequence of similar YGCY motif distribution (Fig 2d-e). This assessment of a limited number of endogenous splicing events indicated that (1) MBNL1 dose-dependent splicing regulation is preserved across our human and murine cell models, and (2) a wide range of dose-response curves described by a range of quantitative parameters can be observed across both systems.

Interestingly, we observed substantial differences in the MBNL1 dose-response for *INSR/Insr* exon 11 between our murine and human dosing models. MBNL1 strongly promoted exon inclusion in ddHEKs (ΔΨ = 0.58) (Fig 2f), consistent with our minigene results (Fig 1c). However, the opposite effect was observed in the ddMEFs where MBNL1 expression mildly stimulated exon 11 exclusion (ΔΨ = −0.10) (Fig 2f). Sequence analysis of the mouse *Insr* exon 11 downstream intron uncovered a low number of putative MBNL1-binding motifs within 300 nucleotides of exon 11 (Fig 2f), consistent with the observed mild effect of MBNL1 expression and established patterns of MBNL positional dependent AS regulation^17^. Interestingly, while the overall number of YGCY motifs was reduced within the murine upstream intron 11 compared to the corresponding human intron, the UGCAUG RBFOX1 binding motif previously identified as critical for MBNL1-dependent exon inclusion activity (Fig 1f) was preserved. Despite the presence of this *cis*-regulatory element, the contrasting effect of MBNL1 expression within the ddMEF system underscores the complexity of AS captured by these dosing models. Furthermore, these studies highlight the capacity of these companion cell lines to also define differences in MBNL1-dependent AS regulation across variable distributions of putative *cis-*motifs and expression of *trans*-acting factors.

### rMATS analysis of RNAseq libraries identifies a collection of MBNL-regulated alternative splicing events affected by RBFOX1

Following our initial assessment and validation of MBNL dose-dependent splicing regulation within both dual-inducible cell lines, we sought to assess the impacts of RBFOX1 on MBNL1-dependent AS outcomes transcriptome-wide. We generated four RNAseq libraries for each double-dosing cell line under the following conditions: (I) no induced expression of either RBP as a control (Fig 3a grey), (II) high MBNL1 expression in the absence of induced RBFOX1 (Fig 3a, green), (III) high RBFOX1 expression in the absence of induced MBNL1 (Fig 3a, orange), and (IV) maximal induced levels of both splicing factors (Fig 3a, brown). Libraries were then prepped and sequenced and AS outcomes comparing the low protein control library (I) to the high MBNL1 (II), high RBFOX1 (III), or high levels of both splicing factors (IV) in both cell models were determined using rMATs. Selecting to exclusively focus on skipped exon (SE) splicing events, a list of common events between these three pairwise comparisons was compiled to capture a subset of events in which MBNL1-dependent splicing is affected by RBFOX1 expression. We then subjected this compiled subset to further filtering to identify events where Ψ was significantly altered by MBNL1 expression (| IΔΨ_II – I_| ≥ 0.1, FDR < 0.05). This analysis yielded a total of 866 and 287 events in ddHEKs and ddMEFs, respectively, which we deemed *MBNL1-regulated*.

**Figure 3:**
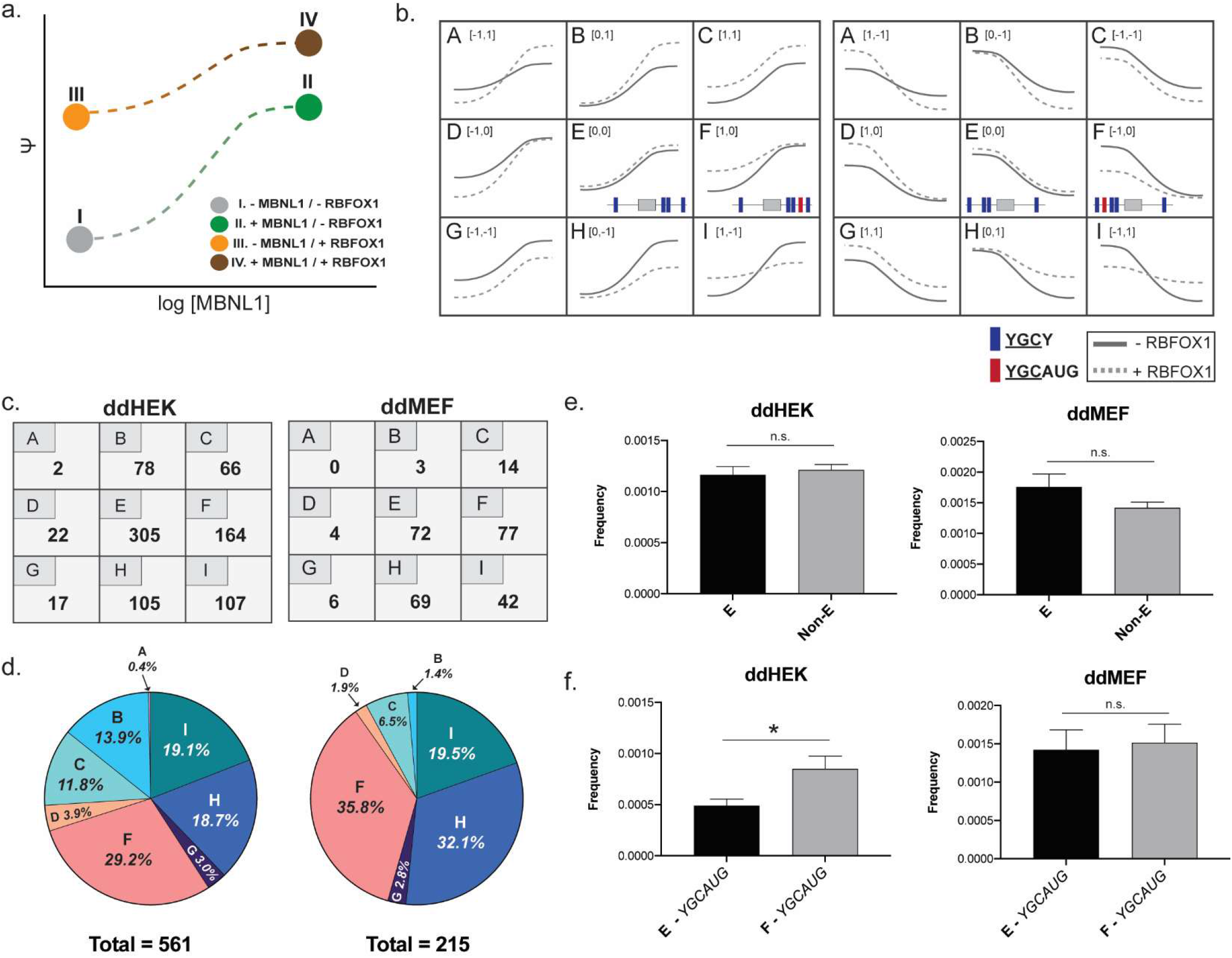
Categorization of RBFOX1 impact on MBNL1 dose-dependent splicing regulation reveals patterns of AS co-regulation transcriptome wide. (a) Four RNAseq libraries were generated (n = 1) for samples with low and high MBNL1 protein concentrations in the presence or absence of RBFOX1 expression within both ddHEKs and ddMEFs. These libraries are representative of the endpoints of MBNL1 dose-response curves (− RBFOX1 and + RBFOX1) as shown by their relative placement on a mock splicing dose-response curve whereby changes in Ψ are plotted against log [MBNL1]. (b) Graphical representation of all potential impacts to MBNL1 dose-response curves based on combinations of positive, negative, or no changes to Ψ at lowest and highest MBNL1 concentrations upon RBFOX1 co-expression. (c) Categorization of MBNL1-regulated events by the effects of RBFOX1 expression on Ψ at low and high MBNL1 expression levels. (d) Pie chart distribution of AS events within each co-regulatory mode (Fig 3c, excluding group E). (e-f) AS events in category E do not show depletion of GCAUG motifs as when compared to all other co-regulatory groups (e). YGCAUG motifs are enriched in co-regulatory mode F compared to group E in ddHEKs, but not ddMEFs (f). Data represented as frequency (total number of motifs/total length of intronic sequence for the 500bp intronic region upstream and downstream of cassette exon) ± SEM, * p < 0.05, n.s. = not significant, unpaired t-test.

### Categorization of effects of RBFOX1 on MBNL1 dose-dependent splicing regulation reveals disproportionate preferences for AS co-regulation

To assess how RBFOX1 co-expression affects MBNL1-regulated AS events we postulated a complete set of mock MBNL1 dose-response curves for which RBFOX1 positively, negatively, or does not significantly alter Ψ at low, high, or both endpoints (Fig 3b). These mock curves are binned into 9 inclusion/exclusion groups (A - I) whereby equivalent impacts on the MBNL1 dose-response for inclusion or exclusion events are placed in matching groups (Fig 3b). To categorize the 866 ddHEK and 287 ddMEF SE events into these 9 groups in an unbiased manner, we devised a binary methodology in which splicing changes that exceeded a significance threshold (ΔΨ_III-I_ / ΔΨ_IV-II_ ≥ 0.1 or ΔΨ_III-I_ / ΔΨ_IV-II_ ≤ −0.1) were assigned “1” or “-1”, respectively, and non-significant changes 0.1 > ΔΨ_III-I_ / ΔΨ_IV-II_ > −0.1) were assigned “0” (Fig 3b, brackets). Each event was assigned to a group based upon the change induced by RBFOX expression at low and high MBNL1 expression. For example, if RBFOX expression resulted in a ΔΨ > −0.1 (scored as −1) at low MBNL expression and a ΔΨ > 0.1 (scored as 1) at high MBNL expression, the event was binned as [−1,1] or group A (Fig 3b).

After all SE events were appropriately binned (Fig 3c) we assessed the general distribution across each co-regulatory group. AS events were in general disproportionately binned across the various postulated groups with a large portion of events in a co-regulatory group (all excluding E). Interestingly in both cell lines, significantly more events in which RBFOX1 exclusively shifted Ψ in the same direction as MBNL1 (groups B, C, F) were identified as compared to those in which RBFOX1 displayed negative co-regulation (groups D, G, H), with twice as many positive co-regulated events observed in the ddHEKs (68.1% vs. 31.9%, respectively). This observation is consistent with established patterns of developmental AS regulation whereby both RBPs have been shown to promote fetal to adult AS transitions^13,29^. When looking at each group individually we found the highest number of events (35% ddHEKs; 25% ddMEFs) bin into group E representing those AS events not impacted by RBFOX1 expression (Fig 3c, d). Of co-regulatory groups, the highest number of events binned into group F (164 in ddHEKs and 77 in ddMEFs), followed by co-regulatory groups H and I (Fig 3c, d). Interestingly, group F most closely resembles the dose-response associated with the *buffering* effects of RBFOX1 observed with the *INSR* minigene (Fig 1). Importantly, this suggests that a comparatively large portion of co-regulated events may utilize a similar shared motif mechanism to facilitate the observed patterns of RBFOX1 and MBNL1 AS co-regulation.

To further support the validity of our binning method and sorting of events into each group, we re-categorized events by increasing the threshold for RBFOX1’s effect on splicing from a ΔΨ value of 0.1 (Fig 3c) to 0.15, 0.2, or 0.3 (Fig S5). As expected, totals for all co-regulatory groups were reduced as the significance cutoff was increased and events were re-categorized into group E (no co-regulation). This reduction in AS events was not as pronounced for group F, especially in the ddHEK system. Group F generally retained high numbers of events (133 events for ddHEKs and 42 for ddMEFs) at the highest threshold (ΔΨ = 0.30, Fig S5c) indicating that most AS events that bin into co-regulatory group F are a consequence of a substantial impact on the MBNL1 dose-response by RBFOX1 expression at low MBNL1 concentrations. Conversely, this observation is consistent with a more subtle effect of RBFOX1 co-expression on other types of co-regulatory modes.

Finally, we assessed conservation based on translated sequence homology for all individual grouped events in both systems. We identified 71 and 74 splicing events of the 866 ddHEK and 287 ddMEF events, respectively, in which the alternatively regulated exonic translated sequence had ≥ 70% identity compared to the matching one-to-one orthologous gene. Of those orthologous events, 16 were binned into matching groups, 9 of which were specific to group F (Table S3). While the total number of orthologous events identified was small, likely due to the distinct species and tissue origin of both cell systems, the large proportion of events with matching patterns of AS co-regulation that binned into group F further emphasizes the prominence of this co-regulatory mechanism. Additionally, these 9 events are good candidates for future targeted mechanistic studies to evaluate the relative importance of specific of *cis*-regulatory motifs that may contribute to the shared patterns of AS outcomes across systems.

Overall, by using postulated mock curves to bin AS events into 9 distinct groups we showed that there is an unequal distribution of events between groups with the majority binned into groups demonstrating co-regulation. Furthermore, in both cell systems the most prevalent amongst those co-regulated (group F) having a similar curve architecture to the INSR minigene experiment suggesting group F highlights an important co-regulatory mechanism in human and mouse.

### Presence of putative RBFOX1 RNA binding motifs does not accurately predict distribution of AS events into specific co-regulatory modes

To assess if the presence or absence of predicted *cis*-regulatory RNA binding motifs explains the distribution of events within AS co-regulation groups (Fig 3c), we examined the 500bp intronic sequence region flanking each skipped exon event for individual motifs. Given that events were initially filtered based on a significant change in Ψ upon MBNL1 expression, we confirmed that >99.5% of events in both systems contained putative YGCY regulatory motifs (data not shown). We first chose to focus on events that bin into group E, which display no co-regulatory capacity. We predicted that these MBNL1-dependent events are unable to be co-regulated by RBFOX1 due to the absence of an RBFOX binding GCAUG motif. Interestingly, analysis revealed that only 40.5% and 25% of events in ddHEKs and ddMEFs, respectively, lacked a GCAUG motif in the flanking intronic regions (data not shown) suggesting a majority of events are in group E not due to the lack of GCAUG motifs. Alternatively, it is possible that the lack of co-regulation is due to an overall relative depletion of GCAUG motifs within the upstream and downstream intronic sequences. To test this alternative hypothesis, we compared the frequency of GCAUG motifs (number of GCAUG motifs/total sequence length) in group E to all other co-regulatory modes. We found that events in group E did not exhibit a significant depletion of GCAUG motifs in either cell system (Fig 3e). This suggests that other factors beyond the presence or absence of GCAUG motifs contribute to lack of response to RBFOX1.

In a similar manner to group E, we next analyzed the distribution of motifs in group F, which has the highest number of binned events of all other co-regulation groups. Furthermore, the dose-response curve architecture of group F most closely resembles the dose response associated with the *buffering* effect of our *INSR* minigene experiment (Fig 1) in which we showed MBNL1 and RBFOX1 use a shared UGCAUG motif to regulate splicing. To assess the potential for other events in group F to also utilize this shared motif mechanism, we scanned the flanking intronic space for the presence of YGCAUG motifs. The leading pyrimidine nucleotide (Y = C or U) allows for an MBNL binding motif (YGCA) to overlap with a RBFOX binding element. Surprisingly, we found that only 30.5% in ddHEKs and 44% in ddMEFs of events in group F contain a YGCAUG motif (data not shown) suggesting a portion of events use a different co-regulatory mechanism. Again, we assessed the alternative hypothesis of whether the frequency of YGCAUG motifs is enriched in this group compared to non co-regulated events (group E). Interestingly, we found that in our ddHEK cell system YGCAUG motifs are in fact significantly enriched in group F compared to E; this enrichment was not observed in the ddMEFs (Fig 3f). This inconsistency is likely due to the significantly lower overall number of SE events (866 ddHEK vs 287 ddMEF, Fig 3c) between our cell models.

Overall, we show here that the absence or presence and distribution of cis-regulatory motifs alone is not sufficient to accurately predict splicing outcomes of individual splicing events for MBNL1 and RBFOX1. Splicing events that are or are not co-regulated by MBNL1 and RBFOX1 exhibit an association beyond that of *cis*-regulatory factors, and both co-regulatory and non-co-regulatory groups are likely dictated by other trans-acting factors or unknown RNA secondary/tertiary structure.

### Full dose-response curves illustrate buffering effects of induced RBFOX1 expression on MBNL1 dose-dependent splicing regulation

To further explore and validate the authenticity of the buffering effects of RBFOX1 expression and *cis*-motif distribution on MBNL1 dose-dependent splicing regulation, endogenous dose-response curves were generated for several splicing events in group F across the complete, inducible MBNL1 concentration range within both dosing models. Based on our experiments with the *INSR* minigene reporter (Fig 1) and ΔΨ thresholds used to categorize co-regulated events, we hypothesized that MBNL1-dependent SE exons within the F module are *buffered* by RBFOX1 due to an overall reduced ΔΨ (|ΔΨ_*−RBFOX1*_| > |ΔΨ_*+RBFOX1*_|) covered over the MBNL1 concentration gradient through shared use of a shared YGCAUG RNA regulatory motif.

*ITGA6*/*Itga6, EXOC1*/*Exoc1*, and *NUMA1*/*Numa1* are examples of three orthologous SE events classified within group F in both ddHEK and ddMEF cell models (Fig 4a-4c). These three events have been previously reported to be regulated by both RBPs, although in an independent context^24,41–45^. Consistent with our RNAseq data, RBFOX1 expression *buffered* the endogenous MBNL1 dose-response by positively shifting exon inclusion at low MBNL1 concentrations with minimal impacts at high MBNL1 levels, leading to an overall reduction in ΔΨ (Table S4). While log (EC_50_) values were not significantly altered, the slope of the dose-response curves in all cases was increased (*ITGA6*/*Itga6, EXOC1*/*Exoc1*) or only minimally decreased (*NUMA1*/*Numa1*), likely as a consequence of the reduced ΔΨ_*+RBFOX1*_ covered over a minimal [MBNL1] range (Table S4). Interestingly, this result indicates an increase in cooperative MBNL1 regulation, potentially as a result of dual use of shared *cis*-regulatory motifs with RBFOX1. In fact, sequence analysis within the flanking regions of these events revealed predicted putative YGCY and GCAUG motifs positionally organized to drive the observed patterns of exon inclusion or exclusion for each RBP (Fig 4a-c). Additionally, as predicted for this co-regulatory mode, the available GCAUG motifs possess a leading pyrimidine base, specifically a <u>U</u>GCAUG, embedded within several putative YGCY motifs. This configuration is predicted, based on our earlier data, to be utilized by both RBPs to facilitate AS co-regulation (Fig 4a-4c).

**Figure 4:**
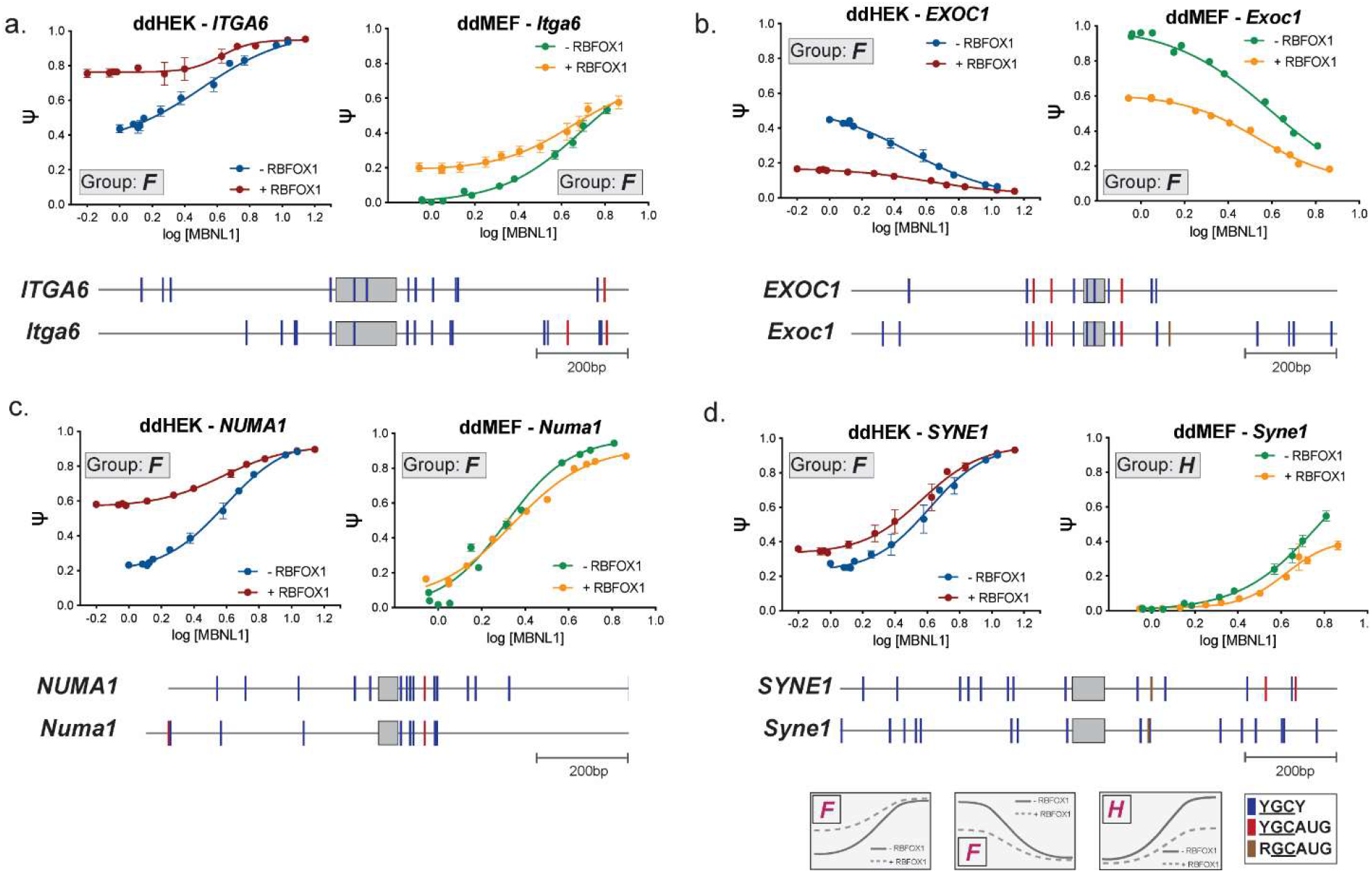
Full splicing dose-response curves illustrate variable impacts of RBFOX1 on MBNL1 dose-dependent splicing regulation. (a-d) MBNL1 dose-response curves in the presence or absence of RBFOX1 expression for ITGA6/Itga6, EXOC1/Exoc1, NUMA1/Numa1, and SYNE1/Syne1, respectively. MBNL1 expression was generated via titration of doxycycline. Additionally, cells were treated with a vehicle control (- RBFOX1) or a single concentration of ponasterone A to induce RBFOX1 expression. Percent exon inclusion (Ψ) was measured for each endogenously expressed mRNA target, plotted against log [MBNL1] levels (Fig 2c), and then fit to a four-parameter dose curve. Data represented as mean ± SEM, n = 3 - 5. The assigned co-regulatory group (A-I) as derived from RNAseq (ΔΨ threshold = 0.10, Fig 3f) is listed for each event and the associated mock curvature displayed below. Quantitative parameters derived from dose-response curves are listed in Table S4. Distribution of putative YGCY, YGCAUG, and RGCAUG binding motifs within and flanking the regulated exon (up to 500 nucleotides) are displayed.

While the distribution of motifs and splicing outcomes was generally preserved for these events within both the ddHEK and ddMEFs (Fig 4a-4c), we observed differential patterns of splicing regulation for SYNE1 in the ddHEK and ddMEF cell lines. *Syne1* displayed a change in the MBNL1 dose-response curve consistent with co-regulatory group H in our ddMEF line but consistent with group F in our ddHEK line (Fig 4d). Analysis of the flanking intronic regions revealed that *SYNE1* contained two additional <u>U</u>GCAUG motifs in the upstream intron in the ddHEK that would be predicted to contribute to the observed buffering effect; these motifs were absent in the corresponding murine intron (Fig 4d). Only a single GCAUG motif was identified with a leading purine nucleotide (R = A or G) predicted to prevent MBNL1 binding while not affecting RBFOX1.

Several other orthologous SE events assayed also did not uniformly sort into the same co-regulatory mode (*NUMB*/*Numb* (E/F), *PLOD2*/*Plod2* (C/F), *ADD3*/*Add3* (F/I), and *INF2*/*Inf2* (D/F) (Fig S6). While some events contained the expected patterns of *cis*-motif distribution to facilitate the observed patterns of exon inclusion across the MBNL1 concentration gradient, no uniform or distinct differences in the distribution of *cis*-regulatory elements adjacent to each cassette exon explained the variability in observed AS co-regulation across systems (Fig S6). Collectively, the endogenous dose-response curves for multiple orthologous SE events underscores the complex variability of AS co-regulation by RBFOX1 and MBNL1 across distinct systems. These differences continue to highlight our lack of understanding of the interplay between both putative *cis*-motifs landscapes and *trans-*acting RBP concentrations and their implications on AS co-regulation.

## Discussion

### RBFOX1 buffers MBNL1 dose-dependent splicing in two distinct cellular systems

In this study we investigated potential modes and mechanisms of AS co-regulation between two splicing factors, MBNL1 and RBFOX1. Using a minigene reporter system and both human and mouse dual-inducible cell lines we evaluated the effects of RBFOX1 expression on MBNL1 dose-dependent splicing regulation. The results demonstrated that MBNL1 and RBFOX1 co-regulate an extensive collection of AS events transcriptome-wide in an interdependent manner. To effectively analyze patterns of AS co-regulation we developed a grouping system to categorize AS events into 9 inclusion/exclusion groups to compare patterns of AS events across cell lines or conditions (Fig 5a). Analysis from our two dual-inducible cell lines revealed that the most predominant co-regulatory group (F) is one in which RBFOX1 *buffers* the MBNL1 dose-response to fine-tune exon inclusion (Fig 3c-d). Based on mutagenesis of the *INSR* exon 11 minigene and in vitro binding experiments, we propose a novel mechanism by which the RBFOX UGCAUG *cis*-regulatory RNA motif is used by both RBPs to co-regulate exon 11 inclusion. More specifically, at low concentrations of MBNL1, RBFOX1 binds to its target UGCAUG binding motif to facilitate exon inclusion. As the concentration of MBNL1 increases, RBFOX1 and MBNL1 compete for the shared site maximizing AS regulation in a non-additive, but cooperative manner (Fig 5b). Via this mechanism, RBFOX1 *buffers* the splicing capacity of MBNL1. Through mutual use of the UGCAUG motif, RBFOX1 and MBNL1 can fine-tune and provide redundancy for AS outcomes, which provides flexibility in maintaining AS patterns beyond that of each RBP alone.

**Figure 5:**
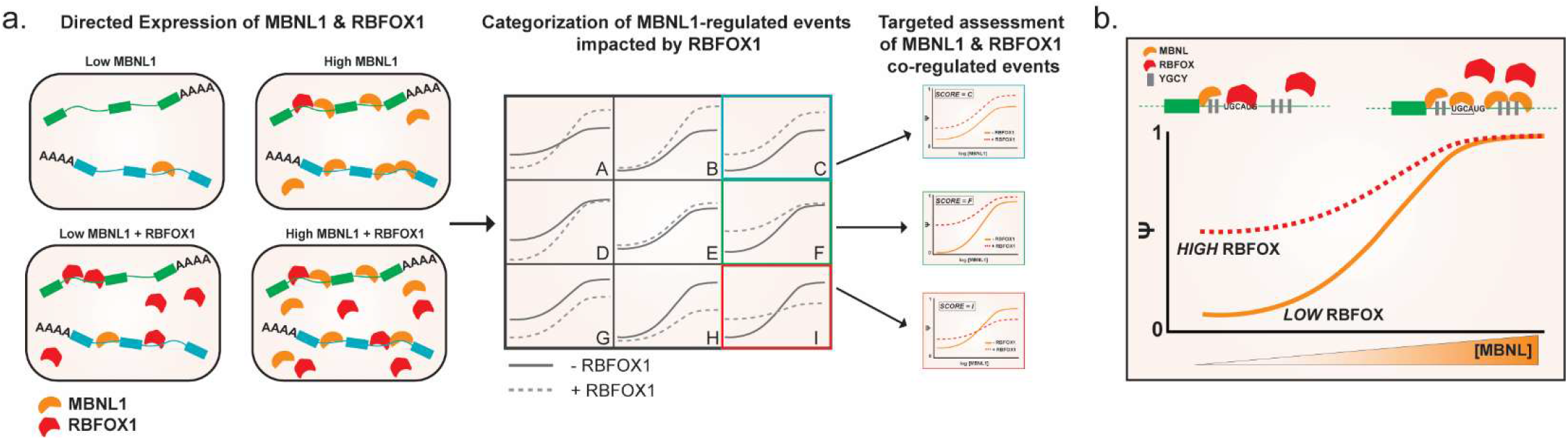
Use of dual-dosing cell models and unbiased categorization of AS co-regulation indicates widespread buffering of MBNL1 dose-dependent splicing by RBFOX1 mediated through use of shared cis-regulatory motifs. (a) RNAseq was performed on dual MBNL1 and RBFOX1 inducible cell lines at minimal and high expression levels of one or both splicing factors. Following identification of AS events regulated by MBNL1 alone, the effects of RBFOX1 expression at the endpoints of the MBNL1 dose-response curve were categorized into 9 predicted co-regulatory modes based on impacts of RBFOX 1 on endpoints of the MBNL1 dose-response curve. (b) Analysis of the impacts of RBFOX1 using the methodology described above in two cell systems revealed the prevalence of co-regulated events in which RBFOX1 buffered MBNL1 concentration dependent-splicing. Experiments with the INSR minigene reporter indicate that this co-regulatory mechanism can be facilitated by shared use the canonical UGCAUG RBFOX binding element. At low concentrations of MBNL, RBFOX binds this cis-regulatory motif to drive exon inclusion. However, at increased levels of MBNL proteins, RBFOX can be displaced and the low-specificity UGCA element is bound by MBNL to maximize exon inclusion.

### Complex interplay of cis-regulatory landscape use by MBNL and RBFOX to facilitate AS outcomes

While MBNL1 and RBFOX1 have previously been shown to bind to the same RNAs using a common motif^31^, here we present evidence that MBNL1 and RBFOX1 compete for use of the same UGCAUG *cis*-regulatory element containing a sub-optimal MBNL binding element (YGCA) (Fig 1). While in the context of the *INSR* minigene, mutation of this site had profound effects on splicing regulation by both RBPs, predicting patterns of global AS co-regulation based on presence of specific *cis*-regulatory elements proved difficult. In fact, minimal RBFOX1 motifs (GCAUG) were not significantly depleted within non co-regulated events (group E) (Fig 3e). Conversely, events that were buffered (group F) by RBFOX1 showed general enrichment for YGCAUG motifs when compared with non-co-regulated events (group E) in ddHEKs, but not ddMEFs (Fig 3f). Furthermore, the general percentage of events in group E that lacked a GCAUG or events in group F that contained a YGCAUG was much lower or higher, respectively, than expected. Collectively, these results suggest that global patterns of AS co-regulation by MBNL1 and RBFOX1 are likely driven by a highly populated and diverse *cis*-regulatory landscape rather than few, high affinity regulatory motifs. Specifically, the expansion of the functional regulatory landscape through use of suboptimal or intermediate affinity motifs may serve to enhance binding to higher-affinity, primary motifs or may only occur at high cellular concentrations to narrow AS activity. Furthermore, additional trans-acting protein factors and/or unknown but biologically relevant RNA secondary or tertiary structure may be important to understanding global patterns of AS. These results speak to the complex use of finite *cis*-regulatory space by a multitude of *trans*-acting factors, especially as it relates to how the regulatory landscape can be altered as concentrations of such factors shift.

### Implications of MBNL1 and RBFOX1 cooperatively regulating AS outcomes in development and disease

Several previous studies have demonstrated cooperative splicing regulation by the RBFOX and MBNL family of RBPs in various systems, including embryonic muscle cell lines^23^, pluripotent stem cell differentiation^24^, and developing mouse cortices^46^. Interestingly, in the context of both stem cell commitment to differentiation and neuronal development, an early induction of RBFOX levels is followed by late-onset increases in MBNL expression. This initial expression of RBFOX proteins facilitates changes in developmental splicing switches that are completed upon MBNL expression, a phenomenon that is mechanistically consistent with our proposed buffering model, in which initial expression of RBFOX1 alters exon inclusion that is maximized as MBNL1 expression increases. Furthermore, many of the events (e.g. *PLOD2*, *ITGA6*, *SYNE2*) previously shown to facilitate developmental maturation via cooperative regulation were also identified and categorized into buffering group F in our systems. Together, these results provide a potential mechanism where differential expression of these two RBPs cooperate during development to interdependently regulate AS transitions.

In addition to MBNL1 and RBFOX1 co-regulating developmental AS transitions^24,46^, more broadly our study highlights the potential for MBNL and RBFOX protein families to functionally compensate for the other to preserve AS patterns in instances where the cellular concentrations or functionality of the other RBP is altered or compromised^31^. This ability may be especially impactful in the context of disease, specifically in the context of Myotonic Dystrophy type 1 (DM1) and type 2 (DM2) in which toxic expansion CUG(G) repeat RNA sequesters and reduces functional MBNL protein levels^47–49^. The presence of RBFOX in MBNL-depleted tissues may compensate for the impacts of functional MBNL loss, potentially mitigating the degree of mis-splicing observed in these tissues. In fact, several events mis-spliced in DM that are associated with disease symptoms are also classified as part of the co-regulatory group F, including *INSR* and *SCN5A*^50–52^. In fact, increased *RBFOX2* expression highly correlates with reduced inferred levels of functional, non-sequestered MBNL proteins based on RNAseq data from a large cohort of DM1 patient muscle^53^. Previous work has also shown DM1/DM2 induced pluripotent stem cell derived cardiomyocytes have increased RBFOX1 protein levels^54^. These results suggest that RBFOX expression may be upregulated in DM as a compensatory mechanism to mitigate disease-specific mis-splicing. As such, changes in expression of the family of RBFOX splicing factors and its impacts on MBNL-regulated splicing events may in part explain phenotypic variation observed in DM patients^55^.

Furthermore, this functional compensation may have important consequences on DM splicing biomarker selection for therapeutic trials. Given the close association between mis-splicing and DM phenotypes, one of the goals of the DM field is the development of effective biomarkers that estimate changes in functional levels of MBNL1 upon therapeutic intervention^18,56^. Naturally, many of these studies focus on MBNL1-dependent AS events as candidate biomarkers. Candidate AS biomarkers that are buffered by RBFOX1 expression such that the changes in splicing are no longer significantly altered by progressive changes in free MBNL1 levels may no longer serve as adequate therapeutic biomarkers. Therefore, it is important to consider RBFOX expression when developing DM biomarkers in model systems.

## Materials and Methods

### Plasmids and cloning

N-terminal HA-tagged RBFOX1 (1-397 amino acids, RNA binding protein fox-1 homolog isoform 4; NCBI accession number NP_061193.2) was synthesized (GenScript, Piscataway, NJ, USA) and cloned into pcDNA3.1+ (Life Technologies, Carlsbad, CA, USA) using standard cloning procedures. Mutations to the wild-type *INSR* minigene^35^ were performed using the QuickChange Site-Directed Mutagenesis platform (Agilent Technologies, Santa Clara, CA, USA) and Phusion high-fidelity DNA polymerase (New England Biolabs (NEB), Ipswich, MA, USA).

### Creation of double dosing cell models

The double dosing HEK-293 (ddHEK) cell line was generated via integration of a ponasterone-A inducible, mOrange-tagged RBFOX1 construct into doxycycline-inducible HA-MBNL1 (amino acids 1-382, NP_066368) HEK-293 cells previously generated and described^18^. The In-Fusion cloning system (Takara Bio, Mountain View, CA, USA) was utilized as per the manufacturer’s instructions to clone a N-terminal, mOrange-tagged RBFOX1 construct into PB-PuroPonA, a vector containing PiggyBac Transposon sequences^57^ flanking a PGK-driven puromycin selection cassette and a minimal CMV promoter downstream of a 5x ecdysone/glucocorticoid response (E/GRE) response elements to facilitate ponasterone A (ponA) inducible expression of mOrange-RBFOX1. HEK-293 cells were transfected with 1 μg of PB-PuroPonA-RBFOX1, 1 μg of a PB-pERV3 (vector containing PiggyBac Transposon sequences^57^ flanking retinoid-X-receptor (RxR) and synthetic VP16-glucocorticoid/ecdysone receptor (VgEcR) CMV promoter-driven expression elements and a puromycin selection cassette), and 1 μg of PiggyBac transposase expression vector (total = 3 μg) using TransIT-LT1 (Mirus Bio, Madison, WI, USA) as per the manufacturer’s instructions. After 24 hours the cells were subjected to puromycin selection (4 μg/mL) (Gibco, Gaithersburg, MD, USA), allowed to recover for several days, and then treated with 10 μM ponasterone-A (Thermo Fisher Scientific, Waltham, MA, USA) for 24 hours. Cells were then sorted for high mOrange expression using FACSAria II (BD BioSciences, San Jose, CA, USA). Individual clones were isolated and the populations expanded in the presence of puromycin (2 μg/mL). A clonal population was selected for experimental use based on GFP-MBNL1 and mOrange-RBFOX1 expression across a range of doxycycline (Sigma Aldrich, St. Louis, MO, USA) and ponasterone A concentrations, respectively.

The double-dosing MEF (ddMEF) cell line was generated via sequential integration of a doxycycline-inducible, GFP-tagged MBNL1 (NP_066368) and ponA-inducible, mOrange-tagged RBFOX1 (NP_061193.2) in *Mbnl1*^−/−^ / *Mbnl2*^−/−^ mouse embryonic fibroblasts (MEFs) gifted by Maurice Swanson. The In-Fusion cloning system (Takara Bio, Mountain View, CA, USA) was utilized as per the manufacturer’s instructions to clone a N-terminal, GFP-tagged MBNL1 expression construct into PB-PuroTet, a vector containing PiggyBac Transposon sequences^57^ flanking a PGK-driven puromycin selection cassette and a minimal CMV promoter downstream of a tetR response element to facilitate doxycycline-inducible expression of GFP-MBNL1. MEFs were initially transfected with 1 μg of PB-PuroTet-MBNL1, 1 μg of PB-Tet-On Advanced (containing PiggyBac Transposon sequences^57^ flanking rtTA-Advanced (Takara Bio) and puromycin selection cassette), and 1 μg of PiggyBac transposase expression vector (total = 3 μg) using TransIT-LT1 (Mirus Bio) as per the manufacturer’s instructions. After 24 hours the cells were subjected to puromycin selection (4 μg/mL), allowed to recover for several days, and then exposed to 1000 ng/mL doxycycline (Sigma Aldrich) for 24 hours. Cells were then sorted for high GFP expression using FACSAria II cell sorter (BD Biosciences). Individual clones were isolated and the populations expanded in the presence of puromycin (2 μg/mL). A single clonal population with high GFP-MBNL1 expression was then transfected as described above to achieve integration of the mOrange-RBFOX1 expression construct. A clonal population was selected for experimental use based on GFP-MBNL1 and mOrange-RBFOX1 expression across a range of doxycycline and ponasterone A concentrations, respectively.

### Cell culture and transfection

HA-MBNL1 HEK-293 cells were routinely cultured as a monolayer in Dulbecco’s modified Eagle’s medium (DMEM) (Corning, Corning, NY, USA) supplemented with 10 % fetal bovine serum (FBS, Corning), 10 μg/mL blasticidin (Gibco), and 150 μg/mL hygromycin (Gibco) at 37 °C under 5 % CO_2_. For all minigene work, cells were plated in twenty-four well plates at a density of 1.5 × 10^5^cells/well. Cells were transfected 24 hours later at approximately 80% confluence. Plasmids were transfected into each well with 1.5 μl of TransIT-293 (Mirus Bio) as per the manufacturer’s protocol. 250 ng of empty vector (pcDNA3.1+, mock) or the pcDNA3.1+ HA-RBFOX1 expression vector were co-transfected with 250ng of a single minigene reporter (total = 500 ng/well). Fresh doxycycline (Sigma Aldrich) was prepared at 1 mg/mL, diluted, and added to the cells at the appropriate concentrations four hours post transfection (10ng/mL). 24 hours post-transfection cells were harvested for experimental analysis.

ddHEKs were routinely cultured as a monolayer in DMEM (Corning) supplemented with 10 % FBS (Corning), 10 μg/mL blasticidin (Gibco), 150 μg/mL hygromycin (Gibco), and 2 μg/mL puromycin (Gibco) at 37 °C under 5 % CO_2_. ddMEFs were also cultured as a monolayer in DMEM (Corning) supplemented with 10% FBS (Corning), 1% penicillin-streptomycin (Gibco), and 2 μg/mL puromycin (Gibco). For experimental setup, cells were plated in twelve-well plates at a density of 6 × 10^4^ cells/well or 3 × 10^5^ cells/well for ddMEFs and ddHEKs, respectively. After 24 hours, fresh doxycycline was prepared at 1 mg/mL, diluted, and then added to the cells at the appropriate concentrations to induce a range of HA-MBNL1 or GFP-MBNL1 protein expression (0 - 10 ng/mL for ddHEKs, 0 - 1000 ng/mL for ddMEFs). To induce mOrange-RBFOX1, fresh ponasterone-A (Thermo Fisher Scientific) was dissolved in 100% ethanol and then added to the media (final concentration = 10 μM). Cells were treated with 100% ethanol as a vehicle control for - RBFOX1 conditions. Cells were then harvested 24 hours post-drug treatment for experimental analysis.

### Cell based splicing assays and curve fitting

Total RNA from HA-MBNL1, doxycycline-inducible HEK-293 cells was isolated using the RNeasy Mini kit (Qiagen, Germantown, MD, USA). The isolated RNA was processed via reverse-transcription (RT)-PCR and Ψ (percent spliced in) for each minigene event as previously described^58^. Total RNA from ddHEKs and ddMEFs were harvested utilizing the Aurum Total RNA Mini kit (Bio-Rad, Hercules, CA, USA) and DNase treated on-column. 1000 ng of RNA was reverse-transcribed using SuperScript IV (Invitrogen, Carlsbad, CA, USA) with random hexamer priming (Integrated DNA Technologies, Coralville, IA, USA) according to the manufacturer’s protocol except that half of the recommended SuperScript IV enzyme was utilized. cDNA was then PCR amplified for 25-32 cycles using flanking exon-specific primers. Primer sequences, annealing temperatures, cycle numbers, and inclusion and exclusion product sizes in base pairs are listed in Supplemental Table 5. PCR-amplified cDNA samples were visualized and quantified using the Fragment Analyzer DNF-905 dsDNA 905 reagent kit, 1-500bp (Agilent Technologies) and associated ProSize data analysis software. Quantified Ψ values were plotted against relative MBNL levels as determined by immunoblot or log (doxycycline (ng/mL)). Curve fitting was performed using the GraphPad Prism software using non-linear curve fitting (log(agonist) vs. response - variable slope (four parameters)) (Ψ = Ψ_min_+ ((Ψ_max_- Ψ_min_) / (1 + 10^((log(EC50) – log[MBNL1]) * slope)^). Where appropriate the minimum and maximum values were restricted to fall between 0 and 1, respectively. Parameters that correlate to biological data, such as log (EC_50_) and slope, were then derived from these curves.

### Protein expression and purification

The RBFOX2 RRM expression vector (XP_016884187, amino acids 100-194) was gifted by Christopher Burge^27^. Briefly, this protein expression vector contains the complete RRM sequence downstream of a GST-SBP tandem affinity tag. The RRM protein was recombinantly expressed using BL21 Star (DE3) cells (Invitrogen) and protein expression induced using 0.5 mM IPTG at an OD_600_ = 0.6-0.7 for 2 hours at 37°C. Following induction, cells were lysed in B-PER bacterial protein extraction reagent (Thermo Fisher Scientific), supplemented with TURBO DNase I (5 U/mL, Thermo Fisher Scientific) and lysozyme (100 mg/mL) for 30 minutes at room temperature. The lysate was then diluted with 1 volume of 1X PBS, incubated on ice for 30 minutes, and centrifuged at 15,000 rpm for 15 minutes. The supernatant was isolated, filtered, and protein loaded onto a GSTrap FF 5 mL column (GE Life Sciences, Marlborough, MA, USA) using the NGC Chromatography system (Bio-RAD). The column was washed twice with 5 column volumes (CV) of 1X PBS followed by 5 CVs of 1X PBS + 1M NaCl. The GST affinity tag was then cleaved using the PreScission Protease (GE Life Sciences) for four hours in cleavage buffer (50 mM Tris pH 7.5, 150 mM NaCl, 1 mM EDTA, 1 mM DTT). The cleaved protein product was then eluted from the column and dialyzed overnight at 4°C into storage buffer (500 mM NaCl, 25 mM Tris pH 7.5, 5 mM BME, 50% glycerol) using a 7000 MWCO Slide-A-Lyzer Dialysis Cassette (Thermo Fisher Scientific). Sample purity was assessed via SDS-PAGE gel and working concentrations determined via Pierce 660nm protein assay reagent and BSA standards (Thermo Fisher Scientific).

N-terminal GST-tagged MBNL1 (amino acids 1-260)^59^ was recombinantly expressed using BL21 Star (DE3) cells (Invitrogen) and protein expression induced using 0.5 mM IPTG at an OD_600_ = 0.6-0.7 for 2 hours at 37°C. Following induction, cells were collected and resuspended in chilled binding buffer (50mM Tris pH 8.0, 500mM NaCl, 1mM DTT, 0.1% Triton X-100). Cells were homogenized using beads (Southern Labware, Cumming, GA, USA) in 4 cycles of 80s homogenization with 3 min in between homogenization. The cell lysate was then centrifuged at 12,000 RPM for 30 min at 4°C and the supernatant isolated and filtered. The filtered lysate was then added to GST-beads (Thermo Fisher Scientific) pre-equilibrated in binding buffer and incubated for 1 hour at 4°C. The beads were then washed 3 times with 20 bead volumes of binding buffer by spinning beads at 750 x g for 3 minutes at 4°C and the supernatant removed. For the third wash, beads were incubated in binding buffer for 15 minutes at 4°C. GST-MBNL1 was eluted from the beads three times by incubating the beads with 2 bead volumes of elution buffer (50mM Tris pH 8.0, 150mM NaCl, pH 8.0, 10mM reduced glutathione) followed by spins as described above. Elutions were combined and dialyzed for 2 hours at 4°C in storage buffer (50mM Tris pH 8.0, 150mM NaCl, 1mM DTT, 50% glycerol) using a 7000 MWCO Slide-A-Lyzer Dialysis Cassette (Thermo Fisher Scientific). Dialyzed GST-MBNL1 was aliquoted, flash frozen in liquid nitrogen, and stored at −80°C. Sample purity was assessed via SDS-PAGE gel and working concentrations determined via Pierce 660nm protein assay reagent and BSA standards (Thermo Fisher Scientific).

### Electrophoretic mobility shift assays

Purified GST-MBNL1 and RBFOX2-RRM proteins as described above were used to conduct the EMSA assay. Fluorescent RNA oligonucleotide probes with a 3’-fluorescein tag were ordered from Integrated DNA Technologies (IDT). Equilibration working solution was made fresh by combining 1mL of equilibration buffer stock (0.01% IGEPAL CA630, 10 mM Tris, pH 8.0, 100 mM NaCl) with 131.6 μL of 100% glycerol and 0.01 mg/mL yeast tRNA (Invitrogen). Serial dilutions of GST-MBNL1 were performed with RNAse/DNase free water on ice. The *INSR* fluorescent RNA probe was heated for 5 min at 60°C and allowed to cool for 2 min. The binding reaction was then assembled in order as follows: 5uL of 10 nM fluorescent RNA probe, 43 μL of equilibration working solution, 2 μL of RBFOX2-RRM (192 nM final concentration), 2 μL of serial dilution GST-MBNL1 and let the binding reaction sit covered from light at room temperature for 30 min. After incubation the binding reaction was loaded onto Criterion 26 well, 10% polyacrylamide TBE precast gels (Bio-Rad) and run in pre-chilled 0.5X TBE (1M Tris, 1M boric ccid, 0.02 M Na2EDTA, pH 8.0) at 100V for 50min. Gels were imaged on ChemiDoc MP (Bio-Rad) and analyzed (Fig 1g) using Bio-Rad Image Lab software.

### Immunoblot

Cells were lysed in RIPA (25 mM Tris-HCl pH 7.6, 150 mM NaCl, 1 % NP-40, 1 % sodium deoxycholate, 0.1 % SDS) (Thermo Fisher Scientific) supplemented with 1 mM phenylmethylsulfonyl fluoride and 1X protease inhibitor cocktail (SigmaFAST, Sigma Aldrich) by light agitation for 15 minutes via vortex. Protein concentrations were assessed via the Pierce BCA assay (ThermoFisher Scientific). 5 μg of lysate was heat denatured for 3 minutes at 95°C for 3 minutes and resolved on a pre-cast 4-15 % SDS-PAGE gels (Bio-Rad) at 200V for 45 minutes in 1X running buffer (25mM Tris pH 8.3, 192 mM glycine, 0.1% (w/v) SDS). The gel was transferred onto a low-fluorescence PVDF membrane (Bio-Rad) for 1 hour at 50V in 1X transfer buffer (25mM Tris pH 8.3, 192mM glycine, 20% methanol (v/v)). The membrane was blocked for 1 hour at room temperature using SeaBlock (ThermoFisher Scientific) and incubated overnight at 4°C with a primary antibody targeting RBFOX1 [1:5000 RBFOX1, ab183348 (Abcam, Cambridge, MA, USA)]. Membranes were then incubated for 1 hour at room temperature with a secondary antibody [1:15,000 donkey anti-mouse 800CW (LI-COR Biosciences, Lincoln, NE, USA)]. Images were acquired using an Odyssey CLx imager (LI-COR). Blots were then stripped using Restore Fluorescent Western Blot Stripping Buffer (Thermo Fisher Scientific) as per the manufacturer’s protocol. Effective antibody removal was assessed via re-imaging of blot. Next, blots were re-blocked in SeaBlock for 1 hour at room temperature and then incubated overnight at 4°C with primary antibodies [1:2,000 MBNL1, 4A8 (Millipore Sigma, Burlington, MA, USA), 1:1,000 GAPDH, 14C10 (Cell Signaling Technology, Danvers, MA, USA)]. Membranes were then incubated for 1 hour at room temperature with secondary antibodies [1:15,000 donkey anti-mouse 800CW (LI-COR), 1:15,000 donkey anti-rabbit 680RD (LI-COR)]. Images were acquired using an Odyssey CLx imager and quantification performed using the associated ImageStudio Lite software (LI-COR). Relative MBNL1 levels in the presence or absence of RBFOX1 co-expression were calculated by first normalizing lanes within the same gel to GAPDH and then by normalizing levels of MBNL1 to the average relative MBNL1 expression level of untreated cells (no MBNL1 or RBFOX1 expression).

### RNAseq Library Construction

Both ddHEK and dMEF cell lines were cultured and dosed following a similar methodology as specified above (see *Cell Culture and Transfection*). 24 hour posed drug treatment, RNA to be utilized for RNAseq was harvested utilizing the Aurum Total RNA Mini kit (Bio-Rad) and DNase treated on column. RNA quality was assessed using the Fragment Analyzer DNF-471 standard sensitivity RNA analysis kit (Agilent Technologies). RNAseq libraries (500 ng, RQN > 9) were prepared using the NEBNext Ultra II Directional RNA Library Kit for Illumina with NEBNext rRNA depletion (Bio-Rad). Protocols were followed as per the manufacturer’s guidelines except that 40X adaptor dilutions were utilized, 4X lower concentrations of index primers were used, all bead incubations were performed at room temperature, and 10 cycles of library amplification were performed. Library size and quality was assessed using the Fragment Analyzer DNF-474 High Sensitivity NGS Fragment Analysis kit (Agilent Technologies). Concentrations of each library were quantified using the KAPA Library Quantification Kit for Illumina (Roche, Basal, Switzerland), pooled, and 2 × 76 paired-end sequencing performed using the Illumina NextSeq 500 (Illumina, San Diego, CA, USA).

### Global splicing analysis

RNA sequencing reads were first cleaned using FASTQC and aligned to the Ensembl reference genomes hg38 (human) and mm10 (mouse) using STAR^60^ with additional flags (--outSAMtype BAM SortedByCoordinate --quantMode GeneCounts --alignEndsType EndToEnd). Splicing analysis was conducted using rMATS (--cstat 0.01 --tstat 4)^61^. Customs scripts were used to further filter exclusively skipped exon events from rMATS output for significant MBNL1-dependent events and to generate a dataset of common events between rMATS pairwise comparisons. Group binning was done manually using the parameters described in the text via Microsoft Excel.

### Conservation

A complete list of human and mouse genes that are one-to-one orthologs was generated using BioMart (release 101). A custom script was used to match one-to-one orthologous genes to events from the 9 splicing groups (Fig 3c) based on the Ensembl gene identifier and access the exonic sequence. NCBI blast toolkit (makeblastdb, tblastx) was used to generate a list of translated sequence homology matches from above events. Custom script was used to remove matches that did not have % translated sequence homology >= 70%

### Custom Scripts

Python scripts used to filter rMATS output, assess conservation, and find motifs are available on GitHub at https://github.com/joeellis1331/Hale_and_Ellis_etal_Coregulation

## Acknowledgements

The authors thank Dr. Maurice Swanson for gift of *Mbnl1* ^−/−^/*Mbnl2* ^−/−^ mouse embryonic fibroblasts and Dr. Christopher Burge for the gift of the RBFOX2-RRM expression vector. The authors would also like to thank Dr. John Cleary and the Berglund Lab for thoughtful discussions and edits to this manuscript.

---

**Supplemental Figure 1:**
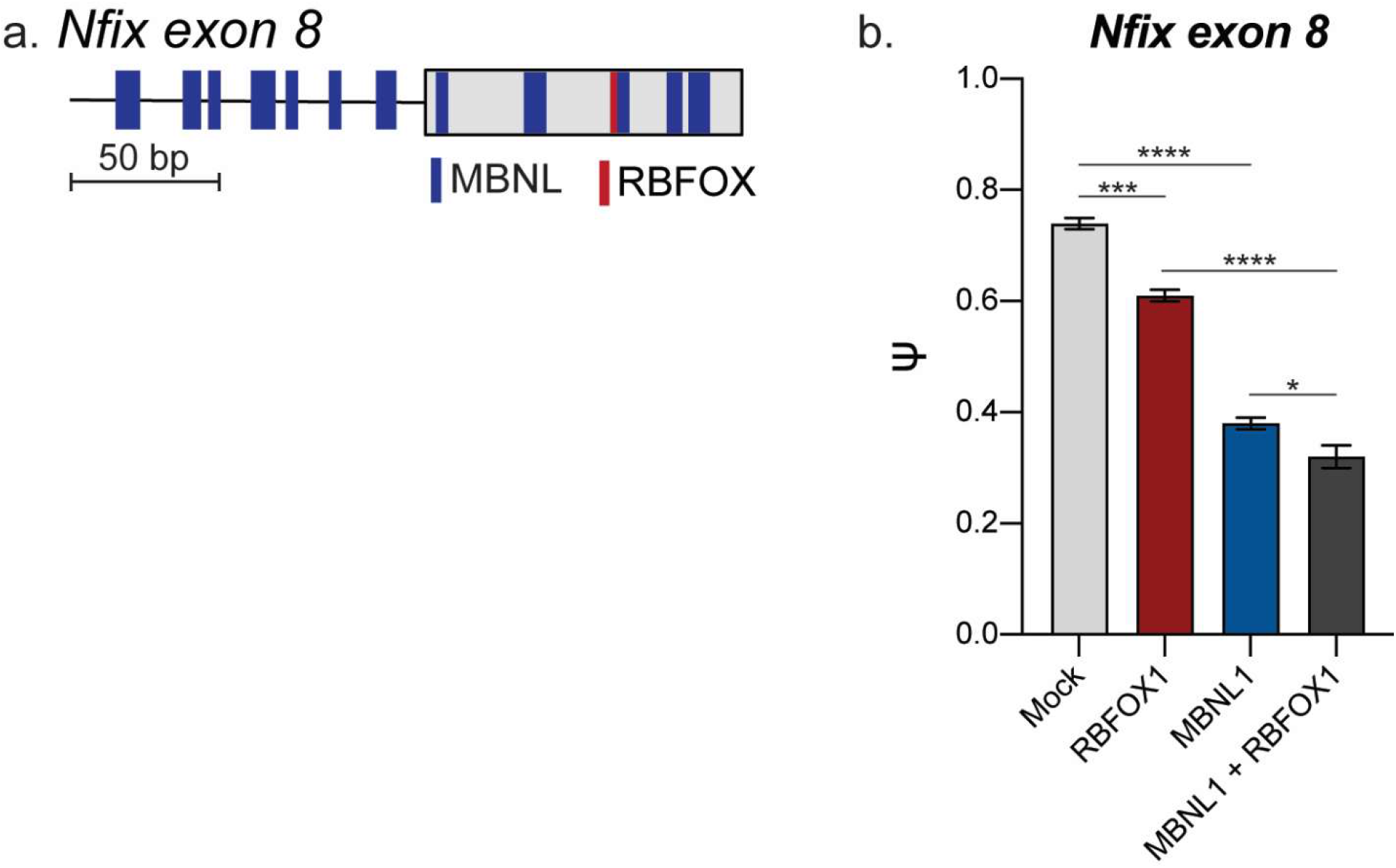
Overexpression of RBFOX1 modestly alters exon inclusion of Nfix exon 8 minigene when co-expressed with HA-MBNL1. (a) Distribution of MBNL1 and RBFOX1 binding motifs in *Nfix* exon 11 minigene. (b) Bar-plot representation of cell-based splicing assay with *Nfix* exon 8 minigene in HA-MBNL1 tetracycline-inducible HEK-293 cells^18^ expressing either HA-MBNL1 or HA-RBFOX1 individually or in combination. Cells were transfected with the minigene reporter and an empty vector (mock) or HA-RBFOX1 expression vector. HA-MBNL1 expression was induced via doxycycline (10 ng/mL). Data represented as mean percent exon inclusion (Ψ) ± SEM, n = 3. (* p < 0.05, *** p < 0.001, **** p < 0.0001, one-way ANOVA).

**Supplemental Figure 2:**
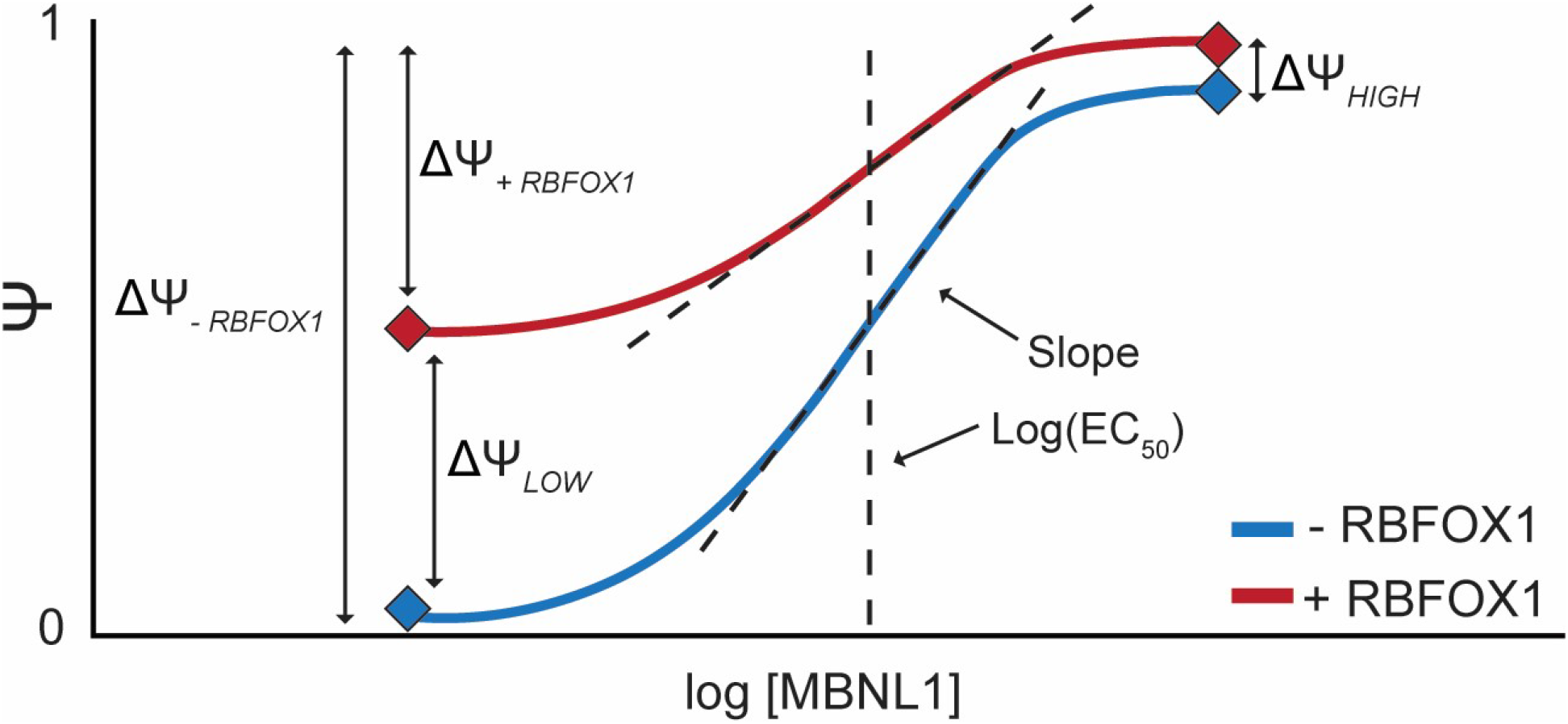
Schematic representation of theoretical change in MBNL1 dose-response curve upon RBFOX1 co-expression. Schematic showing mock MBNL1 dose-response curves displaying theoretical changes in the presence or absence of RBFOX1 expression. Percent exon inclusion (Ψ) is plotted against log [MBNL1] and fit to a four-parameter dose curve. Curve fitting parameters including EC_50_, slope, and span (ΔΨ) are illustrated. In this context, ΔΨ is the difference between minimum and maximum Ψ for each experimental condition (ΔΨ_*- RBFOX1*_ and ΔΨ_+ *RBFOX1*_). Additionally, we illustrate calculation of changes in Ψ at low or high MBNL1 expression levels (ΔΨ_*LOW*_ and ΔΨ_*HIGH*_, respectively). These values are calculated as the difference between the measured minimum Ψ between + RBFOX1 and - RBFOX1 conditions (ΔΨ_*LOW*_) or conversely, the maximum measured Ψ between these same experimental conditions (ΔΨ_*HIGH*_).

**Supplemental Figure 3:**
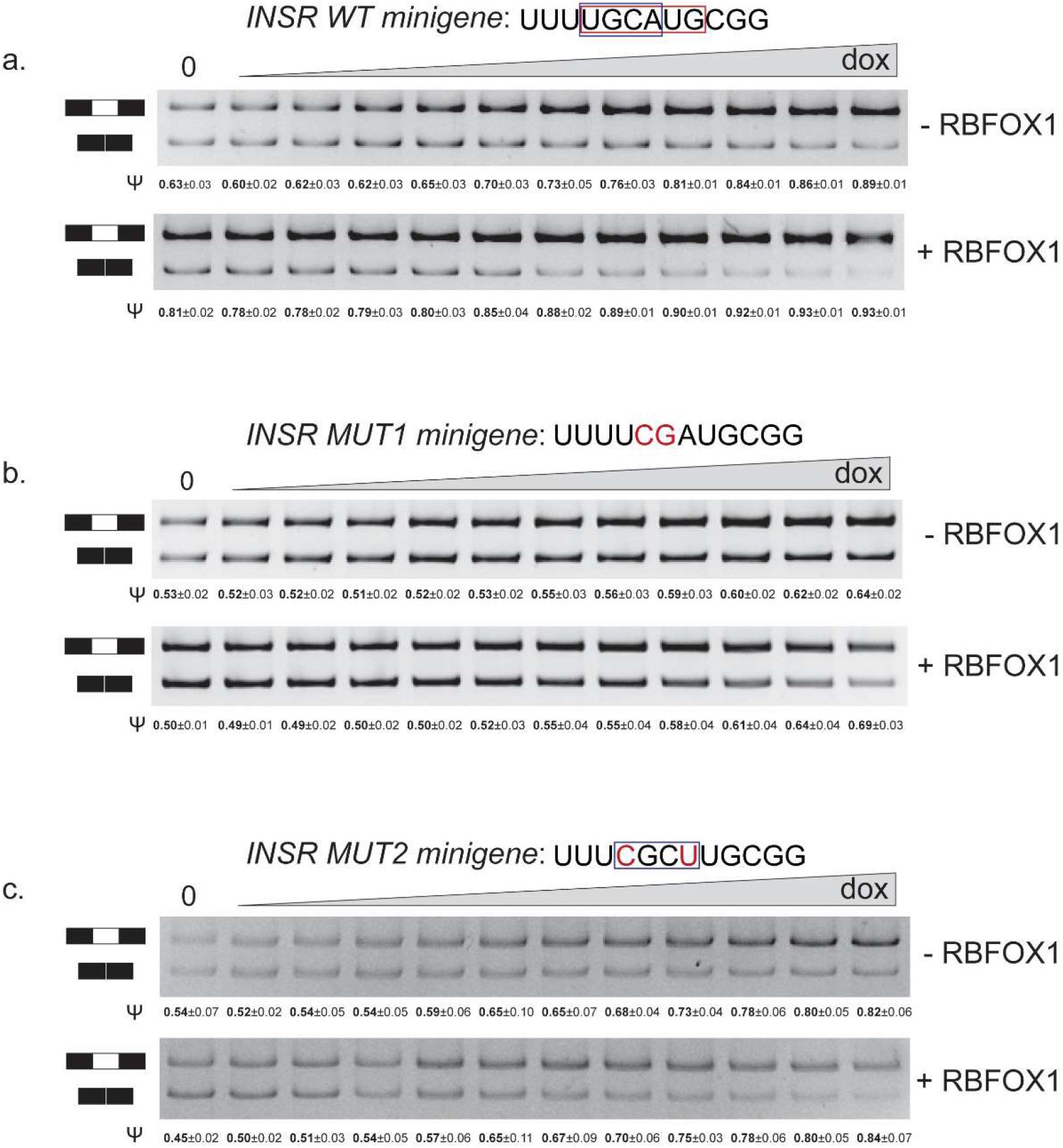
Representative splicing gels for cell-based splicing assays using INSR exon 11 wild-type and mutant minigene reporters in MBNL1-inducible HEK-293 cells. (a-c) Representative splicing gels for wild-type *INSR* minigene, *INSR* MUT1, and *INSR* MUT2 cell-based splicing assays displayed in Fig 1c-e. The mean percent exon 11 inclusion (Ψ) ± SEM (n = 3) for each MBNL1 dose in the presence or absence of RBFOX1 expression is listed below the appropriate lane on each gel. Sequence of minigene containing mutated nucleotides (bolded red) are also displayed and putative RBFOX1 and MBNL1 binding motifs (UGCAUG and YGCY) are boxed in red and blue, respectively.

**Supplemental Figure 4:**
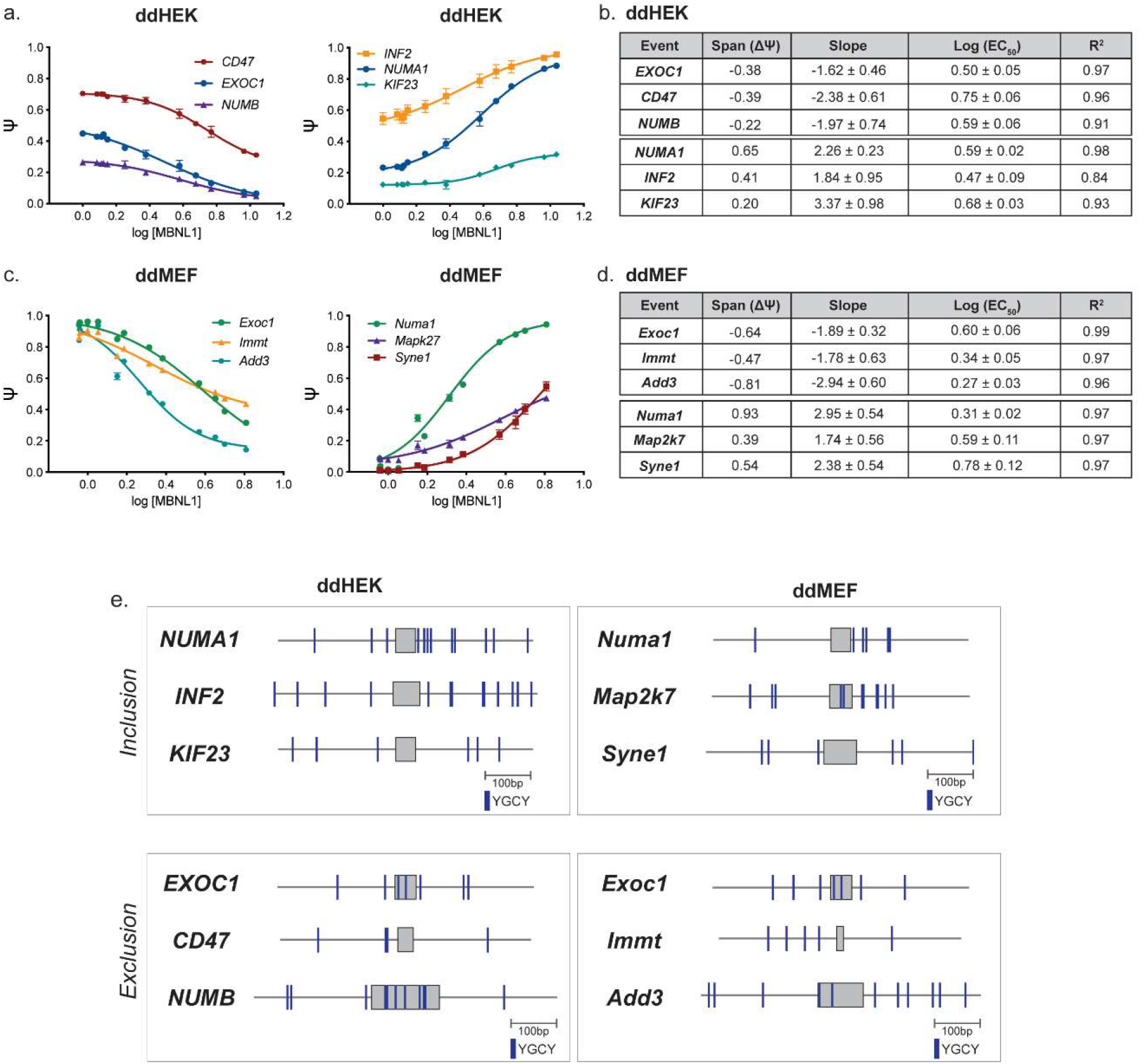
Additional MBNL1 dose-response curves in ddHEKs and ddMEFs and distribution of MBNL1-binding motifs for targeted events. (a-b) MBNL1 dose-response curves for endogenous cassette exon events in ddHEKs and ddMEFs in which MBNL1 expression promotes exon exclusion or inclusion, respectively. Data represented as mean percent exon inclusion (Ψ) ± SEM, n = 3 - 5. (b) Quantitative parameters derived from MBNL1 dose-response curves in (a) and (b), including span (ΔΨ), log (EC_50_), and slope. Data represented as mean ± standard error. (c) Distribution of YGCY motifs within 250bp upstream and downstream of MBNL1-regulated cassette exons for dose-response curves in (a) and (b).

**Supplemental Figure 5:**
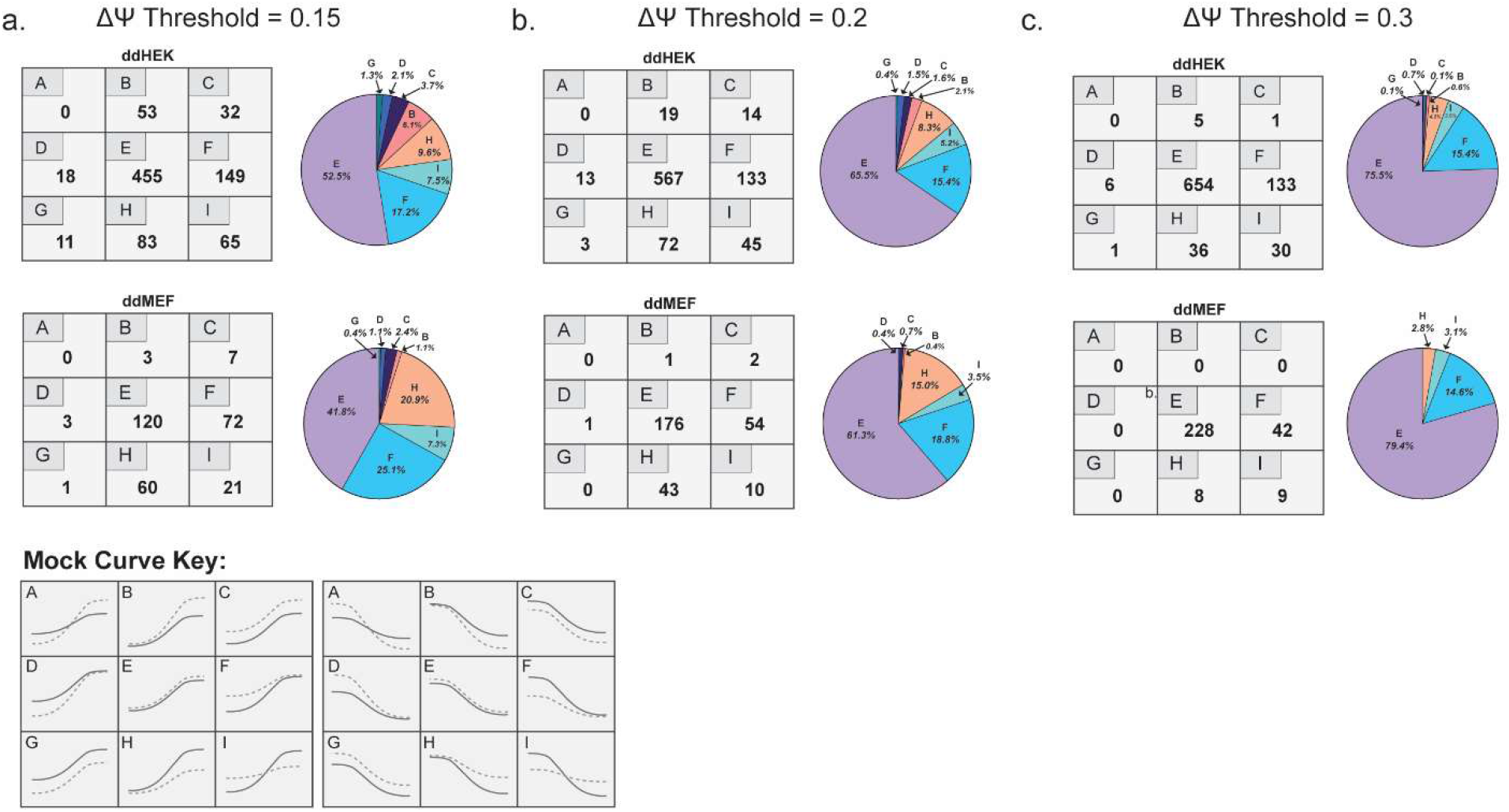
Increasing stringency of significance threshold for categorization of effect of RBFOX1 expression on MBNL1 dose-response does not diminish the number of events within specific co-regulatory modes. (a-c) Categorization of 866 and 287 MBNL1-regulated, SE events in ddHEKs and ddMEFs, respectively (|ΔΨ_II-I_| ≥ 0.1, FDR < 0.05) within increasing significance thresholds of 0.15, 0.2, and 0.3 to define the effects of RBFOX1 co-expression as described in Fig 3. For the purposes of clarity all events regardless of MBNL1 driven inclusion or exclusion activity were binned together and the total number of events within each co-regulatory mode (A-I) is listed. Pie charts showing the percentage of total events assigned to each co-regulatory group as the significance threshold is increased are also displayed next to each matrix.

**Supplemental Figure 6:**
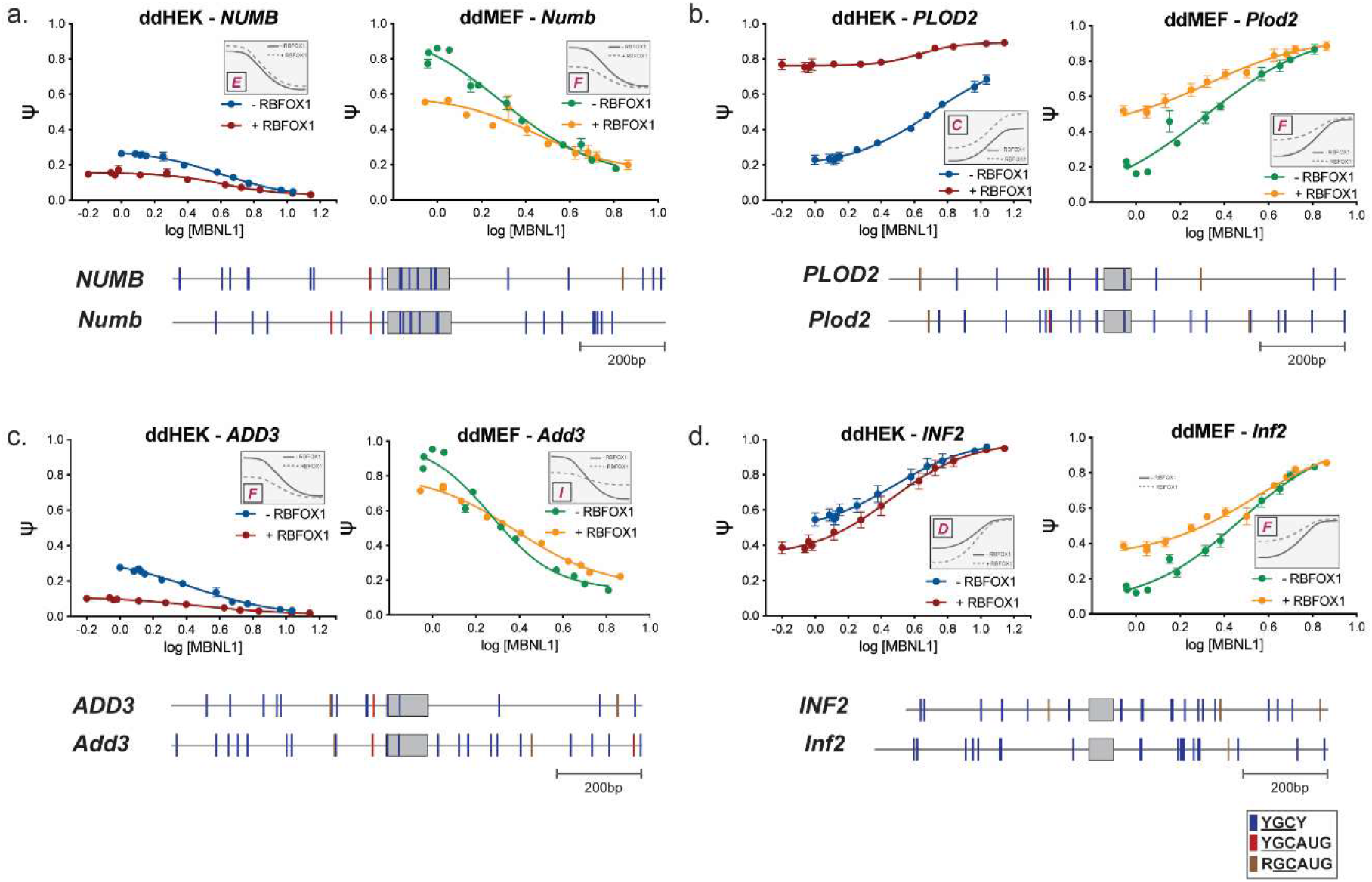
Additional endogenous MBNL1 dose-response curves in the presence or absence of RBFOX1 expression within ddHEK and ddMEF cell systems. (a-d) MBNL1 dose-response curves for orthologous cassette exons *NUMB*/*Numb*, *PLOD2*/*Plod2*, *ADD3*/*Add3*, and *INF2*/*Inf2,* respectively, in the presence or absence of RBFOX1 co-expression. A gradient of MBNL1 expression was generated via titration of doxycycline. Additionally, cells were treated with a vehicle control (- RBFOX1) or a single concentration of ponasterone A to induce RBFOX1 expression. Percent exon inclusion (Ψ) was measured for each endogenously expressed mRNA target, plotted against log [MBNL1] expression levels (Fig 2c), and then fit to a four-parameter dose curve. Data represented as mean Ψ ± SEM, n = 3 - 5. The assigned co-regulatory group (A-I) (ΔΨ threshold = 0.10, Fig 3f) is listed for each event and the associated mock curvature displayed in the inset panel. Quantitative parameters derived from dose-response curves are listed in Table S4. Distribution of putative YGCY, YGCAUG, and RGCAUG binding motifs within and flanking the regulated exon (up to 500 nucleotides) are also shown.

**Supplemental Table 1:** *INSR* Minigene Quantitative Values. Quantitative parameters of MBNL1 dose-response curves for wild-type, MUT1, and MUT2 INSR minigenes in presence or absence of HA-RBFOX1 expression induced via transient transfection in HA-MBNL1 tetracycline-inducible HEK-293 cells. Derived values include span (ΔΨ), slope, and log (EC_50_) in the presence or absence of RBFOX1 expression. Additionally, ΔΨ between - RBFOX1 and + RBFOX1 conditions are listed for the lowest (ΔΨ_*LOW*_) and highest levels (ΔΨ_*HIGH*_) of HA-MBNL1 expression. These values correspond to splicing dose-response curves presented in Fig 1c-e. Data represented as mean ± standard error where appropriate.

| Event | RBFOX1 | Slope | Log (EC<sub>50</sub>) | R<sup>2</sup> | Span (ΔΨ) | ΔΨ<sub>LOW</sub> | ΔΨ<sub>HIGH</sub> |
| --- | --- | --- | --- | --- | --- | --- | --- |
| WT *INSR* | - | 2.27 ± 0.33 | 0.22 ± 0.03 | 0.99 | 0.26 | 0.18 | 0.04 |
|  | + | 3.31 ± 0.67 | 0.13 ± 0.03 | 0.99 | 0.12 |  |  |
| *INSR* MUT1 | - | 2.92 ± 0.60 | 0.37 ± 0.03 | 0.98 | 0.11 | -0.03 | 0.05 |
|  | + | 1.33 ± 0.58 | 0.52 ± 0.13 | 0.98 | 0.19 |  |  |
| *INSR* MUT2 | - | 1.57 ± 0.63 | 0.19 ± 0.10 | 0.97 | 0.28 | -0.09 | 0.02 |
|  | + | 1.91 ± 0.39 | 0.14 ± 0.05 | 0.99 | 0.39 |  |  |

**Supplemental Table 2:** rMATS output with associate co-regulatory groups. Individual sheets for ddHEK and ddMEF data where columns A-O contain extracted rMATS output data containing both inclusion and exclusion SE events and columns P-AC contain co-regulatory group binning of each event using excel formulas using increasing cutoffs (ΔΨ = 0.1, 0.15, 0.2, 0.3)

**Supplemental Table 3:** Orthologous Events and Matching co-regulatory groups. List of human and mouse orthologous splicing events based on one-to-one gene orthologs. Events binned into groups are matched by chromosomal coordinates of upstream exon start and downstream exon end. Cells in rows that are blank contain duplicate events that have been already matched based on start and end exonic coordinates.

**Supplemental Table 4:** Quantitative parameters of MBNL1 dose-response curves for endogenous skipped exon events in presence or absence of RBFOX1 expression in ddHEKs and ddMEFs. Quantitative parameters that describe MBNL1 dose-dependent splicing regulation in presence or absence of induced RBFOX1 expression are listed including slope, log (EC_50_), and span (ΔΨ). Additionally, ΔΨ between - RBFOX1 and + RBFOX1 conditions has been quantified for lowest (ΔΨ_*LOW*_) and the highest (ΔΨ_*HIGH*_) levels of MBNL1 expression (Fig S2). The assigned co-regulatory group categorization (A-I) as derived from RNAseq (ΔΨ threshold = 0.10, Fig 3f) is also listed. Data represented as mean ± standard error.

**ddHEK**
| Event | RBFOX1 | Slope | Log (EC<sub>50</sub>) | R<sup>2</sup> | Span (ΔΨ) | ΔΨ<sub>LOW</sub> | ΔΨ<sub>HIGH</sub> | Co-regulatory Mode |
| --- | --- | --- | --- | --- | --- | --- | --- | --- |
| *ITGA6* | - | 1.65 ± 0.62 | 0.51 ± 0.06 | 0.95 | 0.50 | 0.32 | 0.02 | F |
|  | + | 4.49 ± 2.61 | 0.62 ± 0.06 | 0.73 | 0.20 |  |  |  |
| *EXOC1* | - | -1.62 ± 0.46 | 0.50 ± 0.04 | 0.97 | -0.38 | -0.28 | -0.03 | F |
|  | + | -1.74 ± 0.48 | 0.61 ± 0.06 | 0.94 | -0.13 |  |  |  |
| *NUMA1* | - | 2.26 ± 0.34 | 0.59 ± 0.02 | 0.98 | 0.65 | 0.35 | 0.01 | F |
|  | + | 2.17 ± 0.31 | 0.58 ± 0.03 | 0.98 | 0.32 |  |  |  |
| *NUMB* | - | -1.97 ± 0.74 | 0.59 ± 0.06 | 0.91 | -0.22 | -0.12 | -0.02 | E |
|  | + | -2.46 ± 0.89 | 0.61 ± 0.07 | 0.81 | -0.12 |  |  |  |
| *PLOD2* | - | 1.73 ± 0.60 | 0.72 ± 0.09 | 0.96 | 0.45 | 0.54 | 0.21 | C |
|  | + | 3.56 ± 1.76 | 0.63 ± 0.06 | 0.76 | 0.12 |  |  |  |
| *INF2* | - | 1.84 ± 0.95 | 0.47 ± 0.10 | 0.84 | 0.41 | -0.16 | -0.01 | D |
|  | + | 1.83 ± 0.50 | 0.46 ± 0.06 | 0.92 | 0.56 |  |  |  |
| *SYNE1* | - | 2.50 ± 0.62 | 0.61 ± 0.04 | 0.95 | 0.63 | 0.09 | 0.03 | F |
|  | + | 2.35 ± 0.49 | 0.57 ± 0.04 | 0.95 | 0.57 |  |  |  |
| *ADD3* | - | -1.59 ± 0.55 | 0.43 ± 0.08 | 0.96 | -0.24 | -0.17 | -0.02 | F |
|  | + | -1.62 ± 0.62 | 0.48 ± 0.08 | 0.90 | -0.08 |  |  |  |

| Event | RBFOX1 | Slope | Log (EC<sub>50</sub>) | R<sup>2</sup> | Span (ΔΨ) | ΔΨ<sub>LOW</sub> | ΔΨ<sub>HIGH</sub> | Co-regulatory Mode |
| --- | --- | --- | --- | --- | --- | --- | --- | --- |
| *Itga6* | - | 2.36 ± 0.47 | 0.69 ± 0.08 | 0.97 | 0.53 | 0.20 | 0.04 | F |
|  | + | 2.59 ± 1.01 | 0.65 ± 0.10 | 0.83 | 0.38 |  |  |  |
| *Exoc1* | - | -1.90 ± 0.32 | 0.60 ± 0.06 | 0.99 | -0.64 | -0.37 | -0.13 | F |
|  | + | -2.51 ± 0.26 | 0.54 ± 0.02 | 0.99 | -0.41 |  |  |  |
| *Numa1* | - | 2.95 ± 0.54 | 0.31 ± 0.02 | 0.97 | 0.93 | 0.12 | -0.07 | F |
|  | + | 2.52 ± 0.34 | 0.34 ± 0.02 | 0.98 | 0.72 |  |  |  |
| *Numb* | - | -1.97 ± 0.64 | 0.29 ± 0.05 | 0.95 | -0.68 | -0.29 | 0.02 | F |
|  | + | -2.35 ± 1.09 | 0.45 ± 0.07 | 0.82 | -0.37 |  |  |  |
| *Plod2* | - | 1.74 ± 0.88 | 0.31 ± 0.07 | 0.94 | 0.71 | 0.35 | 0.02 | F |
|  | + | 1.93 ± 0.98 | 0.33 ± 0.08 | 0.89 | 0.38 |  |  |  |
| *Inf2* | - | 2.03 ± 0.55 | 0.47 ± 0.06 | 0.96 | 0.71 | 0.24 | 0.02 | F |
|  | + | 1.94 ± 0.72 | 0.54 ± 0.09 | 0.91 | 0.49 |  |  |  |
| *Syne1* | - | 2.38 ± 0.54 | 0.78 ± 0.12 | 0.97 | 0.54 | 0.004 | -0.17 | H |
|  | + | 4.21 ± 0.99 | 0.62 ± 0.03 | 0.92 | 0.37 |  |  |  |
| *Add3* | - | -2.94 ± 0.60 | 0.27 ± 0.03 | 0.96 | -0.81 | -0.21 | 0.08 | I |
|  | + | -2.13 ± 0.29 | 0.38 ± 0.02 | 0.99 | -0.52 |  |  |  |

**Supplemental Table 5:** Primers utilized for endogenous RT-PCR splicing analysis. Table of primers sequences utilized for evaluation of endogenous exon inclusion levels via RT-PCR analysis in ddHEKs and ddMEFs. PCR annealing temperatures, cycle number, and inclusion/exclusion product sizes (bp) are also listed. Chromosomal coordinates of each skipped exon and flanking exons as identified via rMATS analysis are also provided (hg38 and mm10 for ddHEKs and ddMEFs, respectively).

